# A multiscale approach to investigate fluorescence and NDVI imaging as proxy of photosynthetic traits in wheat

**DOI:** 10.1101/2023.11.10.566533

**Authors:** Nicolas Virlet, João Paulo Pennacchi, Pouria Sadeghi-Tehran, Tom Ashfield, Douglas J. Orr, Elizabete Carmo-Silva, Malcolm J. Hawkesford

## Abstract

With the development of the digital phenotyping, repeated measurements of agronomic traits over time are easily accessible, notably for morphological and phenological traits. However high throughput methods for estimating physiological traits such as photosynthesis are lacking. This study demonstrates the links of fluorescence and reflectance imaging with photosynthetic traits. Two wheat cultivars were grown in pots in a controlled environment. Photosynthesis was characterised by gas-exchange and biochemical analysis at five time points, from booting to 21 days post anthesis. On the same days imaging was performed on the same pots, at leaf and plant scale, using indoor and outdoor phenotyping platforms, respectively. Five image variables (F_v_/F_m_ and NDVI at the whole plant level and F_v_/F_m_, Φ(II)_532_ and Φ(NPQ)_1077_ at the leaf scale) were compared to variables from A-Ci and A-Par curves, biochemical analysis, and fluorescence instruments. The results suggested that the image variables are robust estimators of photosynthetic traits, as long as senescence is driving the variability. Despite contrasting cultivar behaviour, linear regression models which account for the cultivar and the interaction effects, further improved the modelling of photosynthesis indicators. Finally, the results highlight the challenge of discriminating functional to cosmetic stay green genotypes using digital imaging.

**Highlight:** A temporal and multi-scale study of fluorescence and NDVI imaging used as a proxy for photosynthetic parameters

## INTRODUCTION

Modern agriculture is facing multiple challenges to ensure the food security of a growing population. One of them, and not the least, requires tackling the yield gain stagnation in a context of resource-use sustainability and climate change (FAO, 2017). The release of cultivars tolerant to extreme and highly variable climatic conditions is nowadays required (Korres *et al*., 2016). Studies of genotype *vs* environment to improve breeding efficiency and allow rapid varietal adaptation are crucial to avoid yield vulnerability globally and at the farm level (Atlin *et al*., 2017).

In this context, genomics plays a major role, mainly with the identification of genes related to stress tolerance and yield resilience, to inform genomics-assisted breeding for traits related to crop adaptation to climate change (Scheben *et al*., 2016). However, due to the multigenic and complex nature of yield and responses to the environment, many secondary traits, and consequently underpinning genes, will have differential impacts on adaptation in specific environments (Araus *et al*., 2008; Dwivedi *et al*., 2017). In combination with genomics, advances in phenomics are crucial for unravelling the potential of genotypes to adapt in multiple environments (Zhao *et al*., 2019). Currently, genomics is further advanced compared to phenomics, particularly in terms of the development of high-throughput methods and techniques, creating the so-called phenotype-genotype gap (Furbank & Tester, 2011). While plant phenotyping research has surpassed genomics and proteomics fields in terms of the frequency of publications, more developments are necessary to meet the needs of the research community, breeders, and farmers, especially in developing countries where such facilities are less accessible (Costa et al., 2019; Yang et al., 2020). According to Virlet *et al*. (2022), there is uncertainty as to whether the increasing number of phenotypes measured and the depth of information from the new phenotyping platforms will improve the prediction of quantitative traits. Digital HTP provides a non-destructive and non-invasive method for screening large germplasm panels from plant emergence to maturation. HTP often relies on remote sensing technologies to collect information related to morphological, physiological and phenological traits and provide dynamic information on crop development to understand how the yield is built over time and how it is affected by the environment.

Among the parameters required for screening and subsequently selecting the genotypes for yield stability and stress tolerance, physiological traits are of great importance (Großkinsky *et al*., 2015; Shunmugam *et al*., 2018; Sreeman *et al*., 2018). The physiological breeding approach proposes the selection and crossing of parents with different complex, but complementary traits, to achieve cumulative gene action for yield, while selecting progeny using remote sensing, in combination with genomic selection (Reynolds & Langridge, 2016). This is in accordance with the functional phenomics concept, a multidisciplinary approach that includes plant physiology, genomics, phenomics, and bioinformatics, aiming to evaluate ideotypes and generate plant models to predict crop behaviour in different scenarios (York, 2019).

The combination of breeding and high-throughput phenotyping (HTP) of physiological traits in plants can have a significant impact, especially in C3 crops such as wheat. According to the yield potential model proposed by Monteith *et al*., (1977), yield potential is a result of the interaction of the plant with solar radiation through the efficiencies of intercepting and converting light and allocating biomass to the organ of interest, with the conversion efficiency, which is strongly linked to the carbon cycle and photosynthesis, being the variable with highest potential for improvement in C3 plants (Long *et al*., 2015; Zhu *et al*., 2010). Current research focuses on finding proxies for physiological and biochemical parameters in plants, mainly with the advent of remote sensing (Jones & Vaughan, 2010) and the requirement for faster screening of multiple genetic materials (Großkinsky et al., 2015), as well as understanding their interaction with the environment (Pauli et al., 2016). Besides mapping phenological changes and yield-components, the use of these proxies for measuring the efficiency of carbon fixation through photosynthesis has assumed a prominent role in crop sciences (van Bezouw *et al*., 2019). Photosynthesis is generally assessed using gas-exchange instruments, allowing the measurement of the instantaneous variation of CO_2_ assimilation and transpiration. CO_2_ and light response curves (A-C_i_ and A-PAR, respectively) provide access to integrative parameters of the functioning of photosynthesis. From those curves, three primary parameters are generally derived: V_cmax_, J, and A_max_. Derived from A-C_i_ curves, V_cmax_ and J are related to the assimilation rate limited by the available CO_2_ concentration and by the electron transport required for the regeneration of RuBP, respectively. The A-PAR curve provides information about the point where light is no longer a limiting factor for CO_2_ assimilation, often called A_max_ or A_sat_ (Haworth *et al*., 2018). However, collection of CO_2_ and light curve responses as well as net CO_2_ assimilation, is time consuming, and often unfeasible at a population scale in field conditions.

Commonly used techniques for indirectly monitoring photosynthesis are based on spectral data enabling computation of spectral reflectance indices or measuring of sun-induced fluorescence (SIF) based on chlorophyll fluorescence measurement by an active sensor (Porcar-Castell *et al*., 2014; Badgley *et al*., 2017). In general, spectral indices such as NDVI or its derivatives have been used as an indirect measure of primary production, as they are almost linearly correlated to the fraction of absorbed photosynthetically active radiation (Atzberger 2013). However, it has been shown and it is widely accepted that the photochemical reflectance index (PRI) is the only index able to detect short-term variations of CO_2_ assimilation (Dobrowski *et al*., 2005, Merlier *et al*., 2015). Recent studies showed the possibility of predicting V_cmax_, J, and A_max_ combining spectral reflectance and PLSR regression on a small population of wheat in controlled glasshouse conditions and a small set of cultivars in outdoor conditions (Silva-Perez *et al*., 2018). Meacham-Hensold *et al*. (2019) reported that the predictive aspect of the PSLR works across years, but only on the same genetic materials.

The upscaling of those approaches to populations in field conditions is then limited by the need for ground-truth data to build solid and more universal models. The deployment of active sensors for large scale field phenotyping is still complex and so far, few platforms to our knowledge, such as the Field Scanalyzers in the UK and the US, are equipped with chlorophyll fluorescence cameras (Virlet *et al*., 2017; Herritt *et al*., 2020). Laser induced fluorescence transient (LIFT) is another approach which seems more easily usable on field phenotyping trials, but still requires development for large field trials (Wyber *et al*., 2018).

In this study, we compared remote sensing-based reflectance and fluorescence imaging with traditional approaches to measure and evaluate photosynthesis and fluorescence parameters. At the whole plant scale, NDVI and maximum potential quantum yield were imaged under field conditions (Virlet *et al*., 2017). Leaf scale fluorescence imaging was realized in controlled environment conditions to collect maximum potential quantum yield, effective quantum yield of PSII, and non-photochemical quenching parameters (Demidchik *et al*., 2020). Prior to imaging, the same leaves were used for gas exchange and handheld fluorescence measurements, and biochemical analysis were conducted for Rubisco related traits. The biological variability relied on two different wheat cultivars and repeated measurements throughout plant development. Comparison between image variables and traditional approaches was investigated through linear regression models.

## MATERIAL & METHODS

### Plant material and experimental conditions

The plant material was grown in a controlled environment room at Rothamsted Research, Harpenden, UK, from July to October 2018. The plants were grown under a day length of 16 hours, at temperatures of 18/15°C (day/night), relative humidity of 65/75% (day/night) and irradiance of 700 μmols photons m^-2^ s ^-1^ (approximately 500 μmols photons m ^-2^ s ^-1^ at leaf level). Two wheat cultivars (cv. Cadenza and Bobwhite) with contrasting growth and maturation patterns were used. Five wheat seeds were sown in 5 L pots, no thinning was performed, with each pot considered an experimental unit. The experiment was organized into three blocks of five pots of each cultivar, for a total of 30 pots. Zadoks’ scale was used for the evaluation of plant development. Plants were phenotyped at the following developmental stages: booting (Z4.5), anthesis (Z6.5) and 7, 14 and 21 days after anthesis (Z6.5+7, Z6.5+14 and Z6.5+21, respectively). At each time point, one pot per cultivar and per block was used.

### Phenotyping methods

Multiple phenotyping methods were used to evaluate physiological parameters of the plants at whole plant, leaf and biochemical levels. The order of phenotyping was as follows: gas-exchange (A-C_i_ and A-PAR curves), chlorophyll index, light and dark-adapted chlorophyll a fluorescence through handheld devices and the Phenocenter, fluorescence and reflectance through the Field Scanalyzer, and finally, the biochemical analysis.

#### Gas-exchange experiments

The response of net photosynthesis (A) to intercellular CO_2_ concentration (C_i_) and photosynthetic active radiation (PAR) was evaluated through A-C_i_ and A-PAR curves, performed on the central part of flag leaves, using an infra-red gas analyser model LI6400-XT, with an attached leaf chamber LI6400-40 for combined gas-exchange and fluorescence measurements (LI-COR Biosciences). A-C_i_ curves conditions were: flow of 200 µmols s^-1^, block temperature of 20°C, photosynthetic photon flux density (PPFD) of 1800 µmols m^-2^ s^-1^ and relative humidity in the chamber controlled between 55 and 65%. The steps of the A-C_i_ curve were 400, 300, 200, 100, 400, 450, 550, 700, 1000, and 1200 µmols mol ^-1^ for the reference CO_2_ concentration. The A-PAR curves’ conditions for flow, block temperature, and relative humidity were the same for A-C_i_ curves and CO_2_ reference concentration was set at 400 µmols mol^-1^. The steps of the A-PAR curves were 1800, 1400, 1000, 500, 250, 100, 50, and 0 µmols m^-2^ s^-1^ photons.

The data from the A-C_i_ and A-PAR curves were used to calculate additional parameters using the Response Curve Fitting 10.0 and Light Response Curve Fitting 1.0 tools, respectively (available at http://landflux.org/tools). The tool for A-C_i_ curve is based on the parameter calculation method proposed by Ethier and Livingston (2004) while the A-PAR curve toll uses the method developed by Marshall and Biscoe (1980) and Thornley and Johnson (1990). From the A-C_i_ curves, the following parameters were modelled/calculated: maximum carboxylation rate (V_cmax_), CO_2_-saturated electron transport rate (J), chloroplastic CO_2_ concentration (C_c_),inter-cellular CO_2_ concentration (C_i_), intercellular CO_2_ photo-compensation point (C_i_*), chloroplastic CO_2_ photocompensation point (Γ*), CO_2_ compensation point (Γ), CO_2_ transfer conductance (g_i_), mesophyll conductance (g_m_), mitochondrial respiration in the light (R_d_), photorespiration (R_photoresp_), carboxylation efficiency (k_c_^c^). From the A-PAR curves, the following parameters were modelled/calculated: dark respiration (R_dark_), apparent quantum yield (Φ), light compensation point (LCP) and maximum net assimilation rate (A_max_).

#### Fluorescence and SPAD measurements

In addition to the above-mentioned measurements, chlorophyll a content and fluorescence were measured in the same areas of the flag leaves analysed for gas-exchange. The chlorophyll index (SPAD) was measured using the SPAD-502 chlorophyll meter (Konica Minolta). The light-adapted fluorescence parameter, quantum efficiency of PSII (Φ (II) = (F_m_’ -F_s_) / F_m_’) was assessed using the following equipment: Fluorpen FP100 (PSI, Φ (II)FP), FMS2 (Hansatech, Φ (II)FM) and Handy Pea (Hansatech, Φ(II)HP). The dark-adapted fluorescence (F_v_/F_m_HP) was accessed using the Handy Pea after dark adaptation of 30 minutes. Φ(II) was measured with the Fluorpen using the quantum yield, QY function. The saturating light was set up at 3,000 µmols m ^-2^ s^-1^ photons. The FMS2 was used in its default setup (Gain = 50, Mod = 3) and measurement was performed over a 2.5 s period, averaging the equivalent F_s_ signal for 1.8 s before the application of 100-unit saturating pulse over a 0.7 s duration to determine F_m_’. The Handy Pea was used in its default configuration. The OJIP curve was measured under light and dark-adapted conditions with only F_v_/F_m_ considered in both conditions. For the light-adapted leaves, the measured F_v_/F_m_ was assumed to be equivalent to Φ(II).

#### Phenocenter

The same flag leaves that were used for the previous measurements were also used for fluorescence imaging at the Phenocenter (Demidchik *et al*., 2020). The Phenocenter (LemnaTec) is a laboratory-scale, semi-automated system to record and analyse images of plants, through the IMAGING-Pulse-Amplitude-Modulation (PAM) MINI (Waltz). This facility is located in Crop Health and Protection (CHAP), Harpenden, UK. The flag leaves were cut under water using a blade (adapted from Driever *et al*., 2014) and then placed in a water reservoir attached to a black surface using double-sided tape. The tips of the flag leaves were also cut to fit the device and inserted into the Phenocenter (Supplementary Fig. S1). Three flag leaves were analysed at a time. For each leaf sample, three circular areas were defined at 25%, 50%, and 75% of the length of the sample. The three values obtained from each area were averaged to calculate the final sample value for all the measured parameters.

The leaves were left to adapt to darkness for 30 minutes, inside the Phenocenter. Following this, they were exposed to a saturating light pulse, and F_v_/F_m_ was measured (F_v_/F_m_Pc). This measurement was followed by a light curve, with points at each 20 seconds, at the following PAR levels: 1252, 1077, 702, 532, 462, 282, 232, 112, 57, and 2 µmols m ^-2^ s^-1^ photons. At these points the efficiency of the photosystem II (Φ(II)) and the non-photochemical quenching (Φ(NPQ)) were measured. An example of the leaf Φ(II) can be seen on Figure 1.

**Fig. 1.**
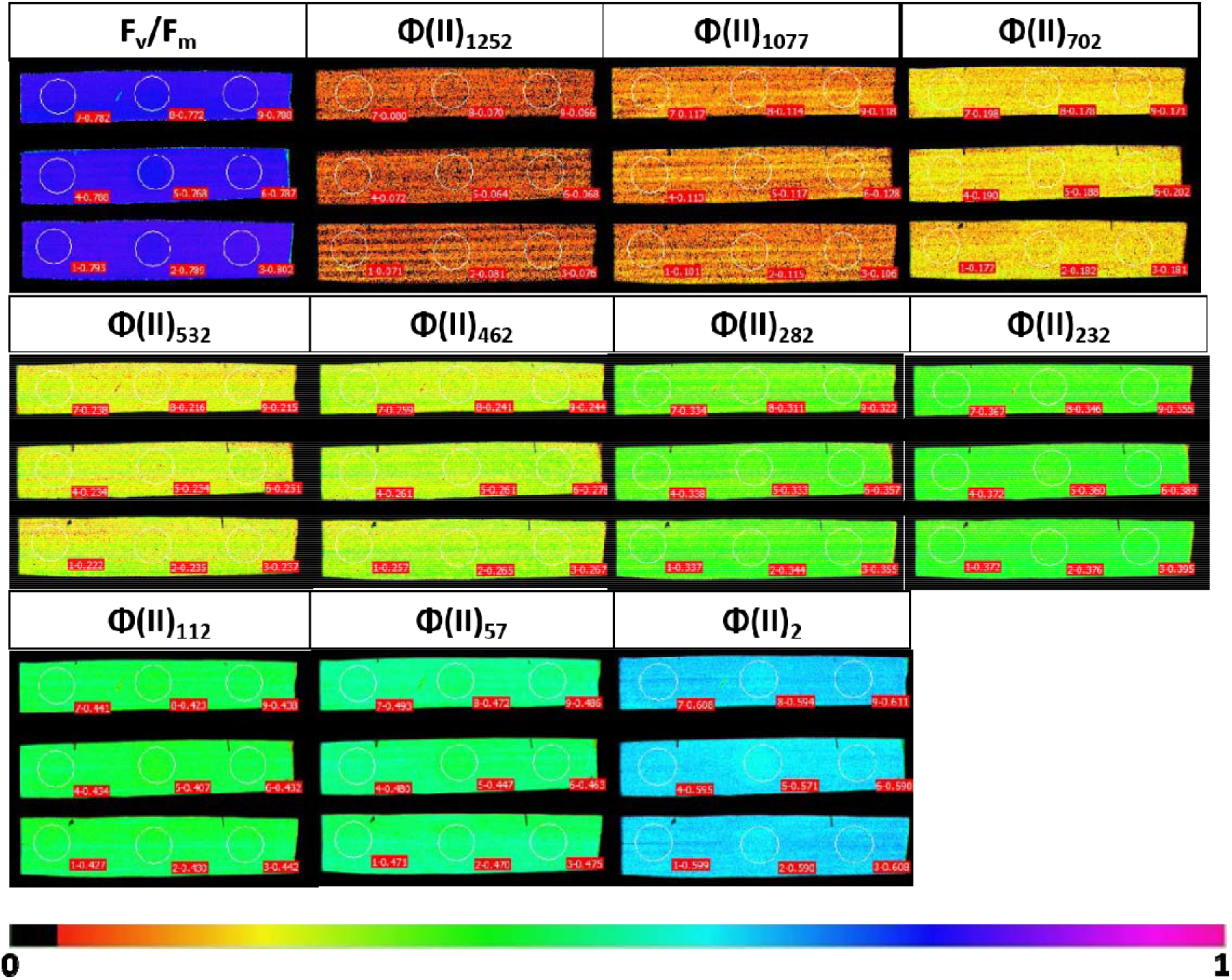
Example Phenocenter images for chlorophyll fluorescence (F_v_/F_m_) and photosystem II quantum efficiency (Φ(II)) analysis. Φ(II)_X_, photosystem II quantum efficiency at the light level of X µmols photons m^-2^ s^-1^.

#### Field Scanalyzer

The pots of the analysed plants were taken to the field and placed on a platform underneath the Field Scanalyzer (LemnaTec) (Supplementary Fig. S2). The Field Scanalyzer is a phenotyping platform (Virlet *et al*., 2017) installed in a field area at Rothamsted Farm, Harpenden, UK. The CropReporter® (Phenovation) (Jalink & van der Schoor, 2011) chlorophyll fluorescence imager and the Hyperspec® Inspector™ VNIR camera (Headwall Photonic) were used to capture images of the entire plants.

The fluorescence was measured at night to collect the F_v_/F_m_ parameter. The camera was positioned at 1m above the top of the plant, and a serie of flashes of red light (620nm) was used to induce the chlorophyll fluorescence (CF) response for 1400 ms (4400 µmols m^−2^ s^−1^ at 70 cm from the bottom of the camera) saturating the electron transfer between the two photosystems (PSII and PSI), and records 24 images within that time. The exposure time of the first image was initiated using the trigger at the rising edge of LED illumination, and this image was called ground fluorescence (F_0_). The image with the maximum fluorescence (F_m_) was then processed to create a mask to remove the background using the Otsu thresholding approach, which was then applied to the F_0_ image. A composite image is then created by combining F_m_ and F_0_ images to obtain the F_v_/F_m_ image to extract the F_v_/F_m_ average values from the plant. Figure 2 shows the final F_v_/F_m_Sc image obtained for Bobwhite and Cadenza at each time point for one of the repetitions.

**Fig. 2.**
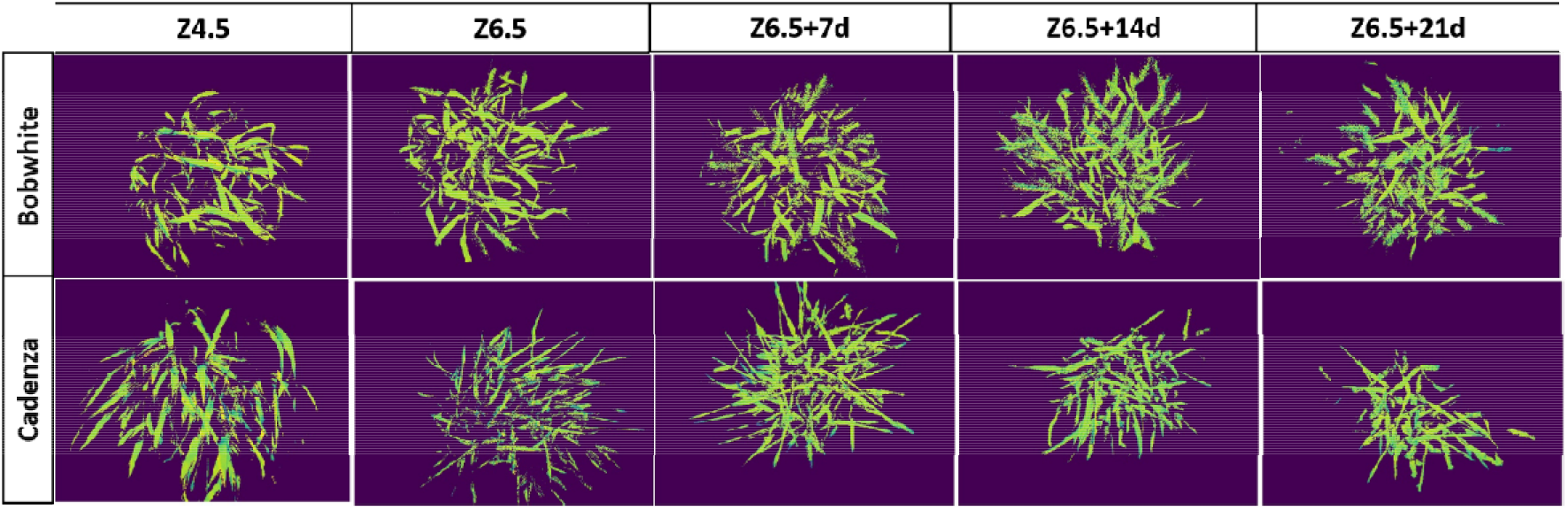
Example Field Scanalyzer images for chlorophyll fluorescence (F_v_/F_m_Sc). Z4.5, booting stage; Z6.5, anthesis; Z6.5 + 7d, 7 days post-anthesis; Z6.5 + 14d, 14 days post-anthesis; Z6.5 + 21d, 21 days post-anthesis.

The hyperspectral images were captured on the same day or the following day of the fluorescence measurement, depending on the weather conditions. A white reference panel, Ultra-lightweight Reflectance Target (Zenith Lite™ Ultralight Targets 95%R, Sphereoptics®), mounted on a tripod was also scanned during the acquisition. This panel was used later to convert the raw data into reflectance data using the method describe by Sadeghi-Tehran *et al*. (2021). After segmenting the plant from the background, NDVI was computed from the 680 and 800 nm wavelengths.

#### Biochemical analysis

The same cut leaves used for the Phenocenter analysis were kept in a tube with water and were transferred to a cabinet to activate photosynthesis and Rubisco for 40 mins before sampling. Cabinet conditions were maintained at 25°C, 70% relative humidity and an irradiance of 900 μmols photons m^-2^ s ^-1^ at the leaf level. The samples were taken from the centre of the area using a freeze clamp with 2 cm head diameter, and sample area calculated by measuring the sample width. The samples were immediately transferred to foil packets, snap frozen in liquid nitrogen, and stored at −80°C until analysis.

Proteins were extracted from leaf samples as described by Carmo-Silva *et al*. (2017). This protein extract was used to determine Rubisco activity via incorporation of ^14^CO_2_ in acid-stable products (Carmo-Silva *et al*., 2017), and activation state was calculated from the ratio of initial/total Rubisco activity (V_i_, V_t_). An aliquot of the homogenate was used to determine chlorophyll a and b content (Chl a and Chl b) using ethanol (Wintermans & de Mots, 1965) and measuring absorbance in a microplate reader SPECTROstar Nano (BMG LabTech). The supernatant prepared for Rubisco activity was also used to quantify total soluble protein (TSP) via the Bradford assay (Bradford, 1976), and the Rubisco content (Rubisco) was determined by [^14^C] carboxyarabinitol-1,5-bisphosphate (^14^C-CABP) binding assay (Whitney *et al*., 1999).

### Data analysis

In order to conduct gas-exchange, fluorescence and SPAD measurements, as well as Phenocenter and biochemical analysis, two plants of each pot were used as technical replicates, and their values were averaged to form one biological replication for each block. However, for the Scanalyzer analysis, only one value was determined for the whole pot.

The data was organized in two factors: two cultivars (cv. Bobwhite and Cadenza) and five developmental stages (Z4.5, Z6.5, Z6.5+7, Z6.5+14 and Z6.5+21), with three blocks, resulting in a total of 30 points. The parameters were split into two groups: image-based parameters obtained from the Field Scanalyzer and the Phenocenter, and the more “traditional” parameters collected by gas-exchange, biochemical analysis and handheld devices.

The relationship between the image-based variables and each of the traditional parameters was investigated using linear regression models. The first approach looked at the relationship between the parameters of the two groups (image-based and traditional methods) for each cultivar individually, but with all the development stages (Individual models for Bobwhite and Cadenza). In the second approach, the linear regression was fitted using the whole dataset to analyse the generalist capacity of the regression and analyse the effect of the cultivars. Two models were built for this purpose: one model considered only the regression between the variables of the two groups without the cultivar effect (Generalist model), and the other model considered the cultivar effect (cv.) and its interaction (Int.) with the considered parameter (Interaction model).

All the results from the first approach are presented in supplementary data (Supplementary Tables S1 to S4). A subsample is presented in the results section. A subset of five image-based variables were selected and compared to six parameters derived from the CO_2_ and light curve response (V_cmax_, J, g_m_, k ^c^, R and A), to six parameters resulting from the biochemical analysis (TSP, Chl a, Chl b, Rubisco content, V_i_ and V_t_) and five parameters collected by the handheld device (SPAD, Φ(II)FP, Φ(II)FM, Φ(II)HP and F_v_/F_m_HP). The image-based variables selected were the fluorescence parameters collected with the Field Scanalyzer (F_v_/F_m_Sc), and with the Phenocenter (F_v_/F_m_Pc, Φ(II)_532_ and Φ(NPQ)_1077_) as well as the NDVI calculated from the reflectance data.

ANOVAS were conducted to analyze the results of the regressions for each arrangement. The results of the F-test of the ANOVA (F and *p*-values) and the adjusted coefficient of the determination (R ^2^_adj_) were used to evaluate the regressions. Tables 1-5 display the F and *p*-values and R ^2^_adj_ for the Individual Models (Bobwhite then Cadenza), the Generalist Models and the Interactions Models. In Interactions Models, the F and *p*-values of the ANOVA for the parameter in the y-axis are referred to as F (Par.) and *p* (Par.), respectively. The cultivar effect F (cv.) and *p* (cv.) and the interaction effect between cultivars F (Int.) and *p* (Int.) are also noted. For *p* (Par.), a value lower than 0.05 suggests a significant linear correlation between the two analysed parameters. A p-value lower than 0.05 for *p* (cv.) and *p* (Int.) indicates that the intercepts and the slopes of the fitted regression for each cultivar are significantly different. The *p* (Model) and R ^2^_adj_ are, respectively, the general *p*-value and the R ^2^_adj_ of the linear regression with the line effect. The cultivar effect tests the hypothesis that the intercept is not different between the two cultivars. A significant effect indicates a statistical difference between the intercept of the two cultivars as well as the two regression lines are mostly parallels. The interaction effect tests the null hypothesis that there is no difference in slopes between the two cultivars, which indicates that both cultivars are responding similarly.

**Table 1.**
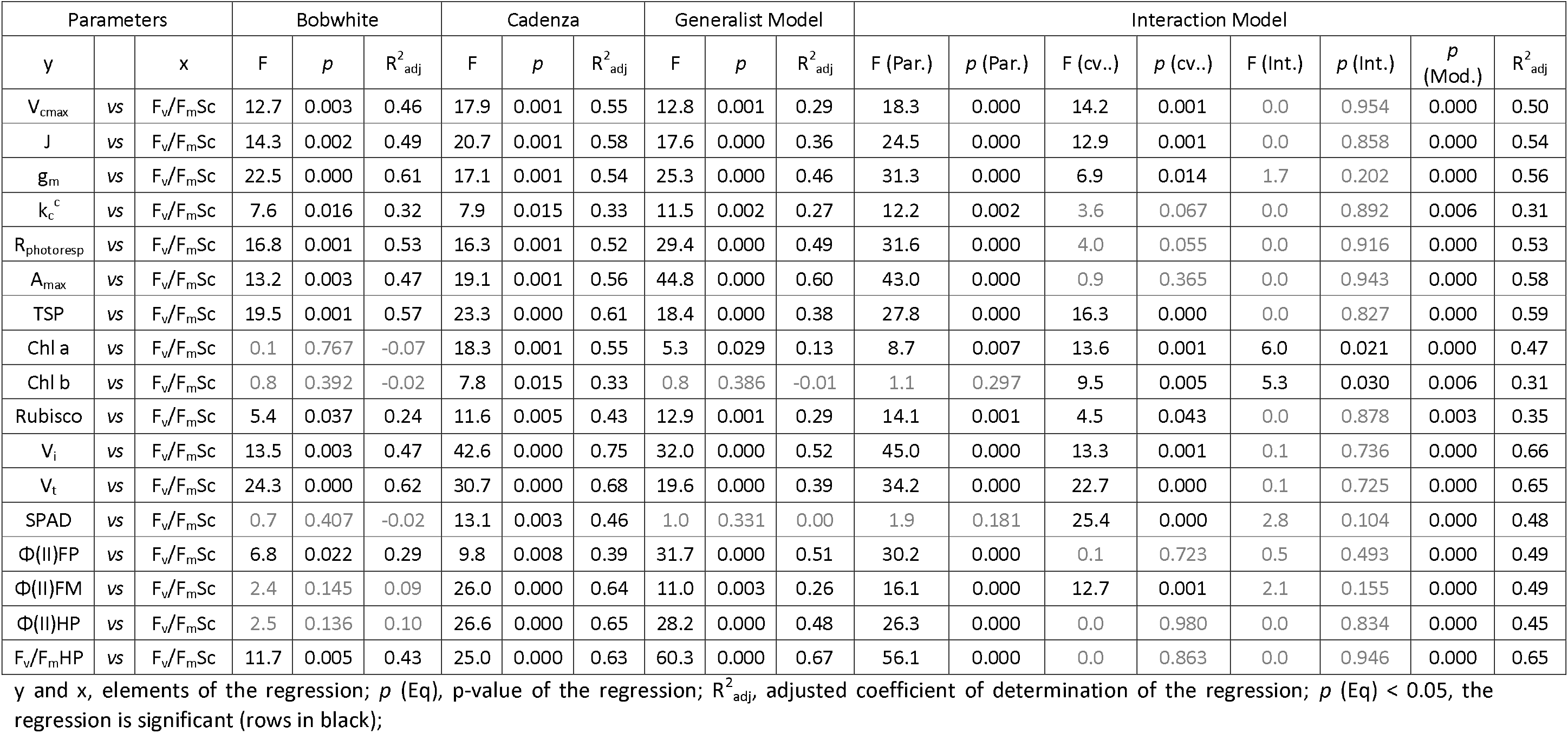
Parameters of linear regressions between chlorophyll fluorescence (F_v_/F_m_) measured by the Field Scanalyzer and other measured parameters.

The residuals of the linear regressions were evaluated for normality, homoscedasticity (homogeneity of variances) and independence by visually checking the Q-Q, residuals *vs* fitted values and histogram of residuals’ plots, and also by the Shapiro-Wilk test. The most relevant linear regressions were plotted (Supplementary Fig. S3-S7) and the results were presented in tables 1 to 5. The R software was used for the analyses with the following packages *ggpmisc*, *stats* and *olsrr* (R Core Team, 2020).

## RESULTS

### Date and cultivar effect

Among the presented variables, two main patterns were observed (Fig. S3, Supplementary Table S5). The first showed a significant decrease of the values for both cultivars from booting (Z4.5) to 21 days post anthesis (Z6.5+21) for all the variables derived from the gas-exchange (V_cmax_, J, g_m_, k_c_^c^, R_photoresp_ and A_max_), for the biochemical parameters related to the Rubisco (TSP, Rubisco, V_i_ and V_t_) as well as for Φ(II)HP and F_v_/F_m_HP. The same was observed for the image variable Φ(NPQ)_1077_. The second pattern showed a contrasted behaviour between the two cultivars for Φ(II)FM and the image variables F_v_/F_m_Sc, F_v_/F_m_Pc, and NDVI with a significant decrease overtime for Cadenza, whilst no significance differences were observed for Bobwhite.

The chlorophyll related parameters (Chl a, Chl b and SPAD) showed an increase of the values peaking at Z6.5+7, followed by a decrease for Cadenza and no significant variation for Bobwhite. For Φ(II)_532_, the values decreased significantly Z6.5+14 for Cadenza, whilst for Bobwhite the values dropped significantly after booting with no more variation for the rest of the experiment. Finally, Φ(II)FP did not show significant variation for either cultivar with, however, a decreasing trend observed for Cadenza at Z6.5+14 and Z6.5+21.

### Field Scanalyzer

#### Dark-adapted chlorophyll fluorescence (F_v_/F_m_Sc)

The F_v_/F_m_Sc calculated from images of the Field Scanalyzer for whole plants (Fig. 2) showed high correlations with most of the parameters measured by traditional methods (Table 1; Supplementary Fig. S3). Cadenza showed significant correlations with all the parameters, with the highest fitting for V_i_ (R ^2^_adj_ = 0.75) and the lowest fitting for k_c_^c^ and Chl b (R ^2^_adj_ = 0.33). For Bobwhite, the regressions between F_v_/F_m_Sc and (i) the gas-exchange parameters were all significant (R ^2^ = 0.32 – 0.61), (ii) the biochemical parameters were mostly all significant (R ^2^_adj_ = 0.24 – 0.62) except for the Chl a and b content and (iii) the handheld devices were significant only for Φ(II)FP (R ^2^_adj_ = 0.29) and F_v_/F_m_HP (R^2^_adj_ = 0.43).

The choice of the cultivar appeared to impact the relationship between the traditional parameters and the imaging F_v_/F_m_Sc (Fig. 3). The Interaction Models (Table 1) confirmed that most of the parameters displayed a significant cultivar (cv.) effect except for the parameters k_c_^c^, R_photoresp,_ A_max_, Φ(II)FP, Φ(II)HP, F_v_/F_m_HP. When the cv. effect was significant, the R ^2^_adj_ from the Interaction Model increased compared to the Generalist Model. On the other hand, the R ^2^_adj_ between both models were quite similar when no significant cv. effect was observed. No significant interaction between the parameters and cultivar effect was observed, except for Chl a and b, indicating that in most cases, there is no significant difference between the slopes of the regression of the parameters and F_v_/F_m_Sc of each cultivar.

**Fig. 3.**
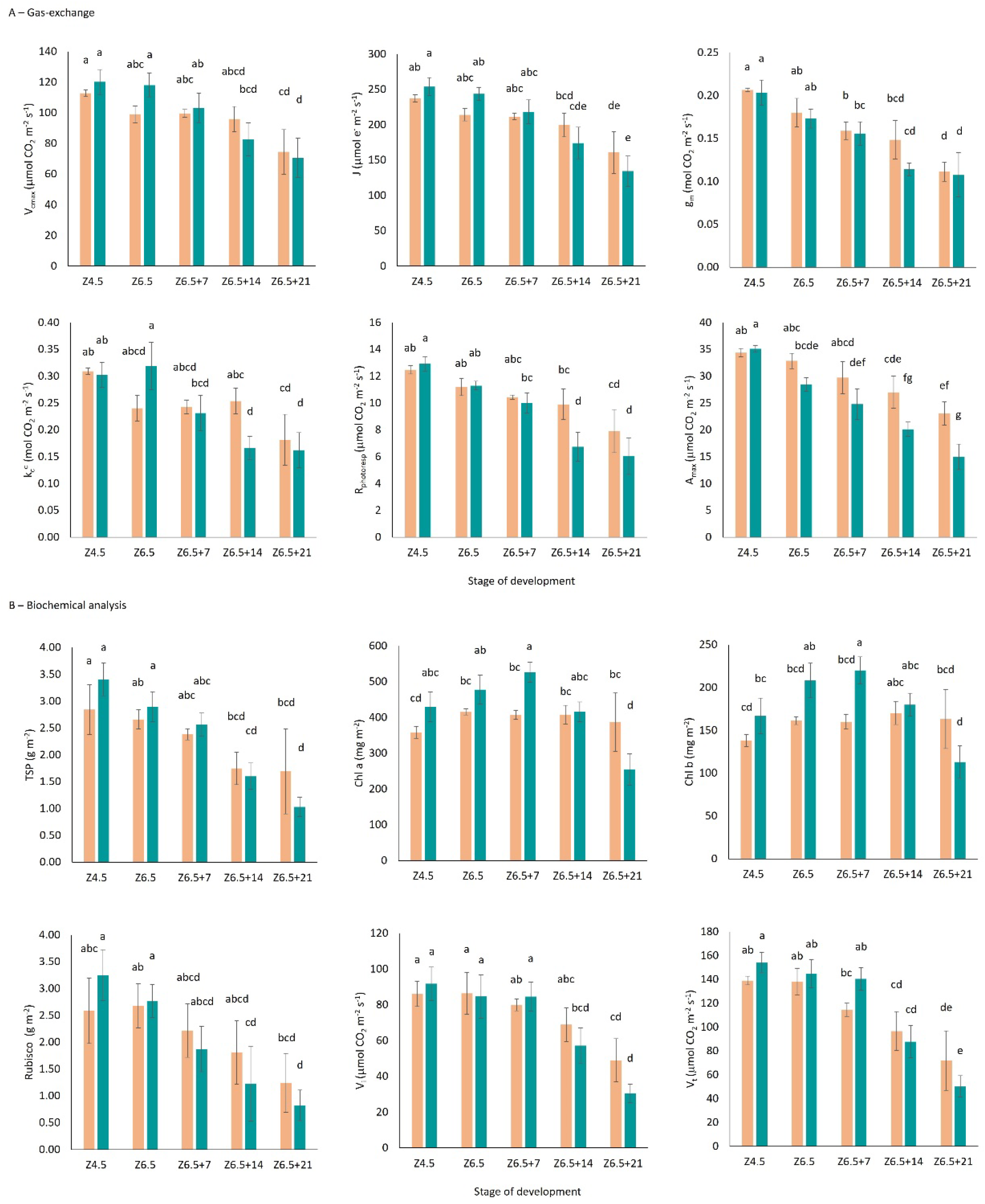
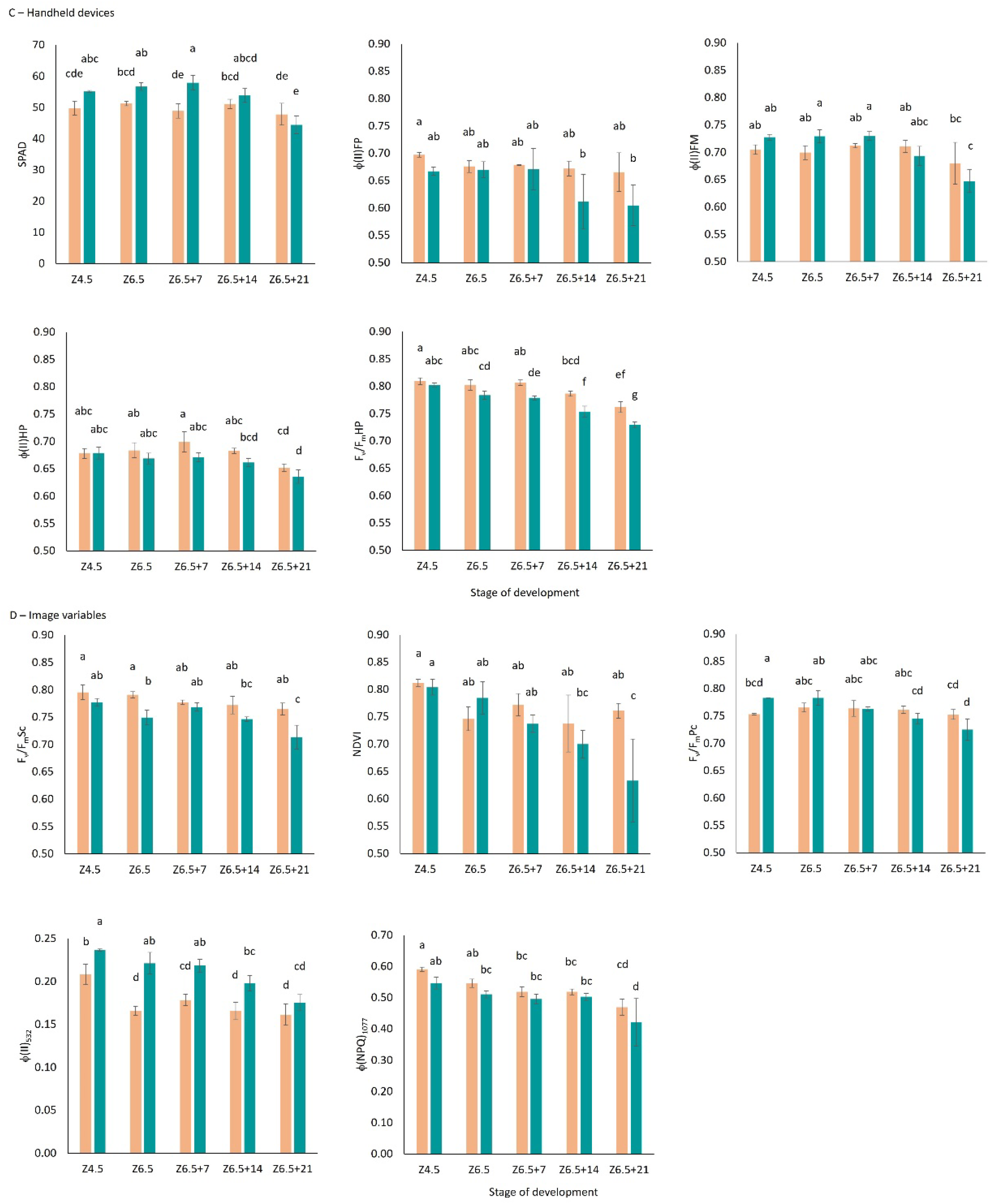
Mean and standard deviation for each variable for Bobwhite (orange) and Cadenza (blue) at booting (Z4.5), anthesis (Z6.5) and at 7-, 14-and 21-days post anthesis (Z6.5+7, +14, +21) for (A) gas-exchange, (B) biochemical, (C), handheld devices and, (D) image variables. Letters come from the post-hoc test *HSD.test* allowing multiple comparisons of treatments (here time point and cultivar) by means of Tukey. Different letters indicate a statistical difference between cultivar and/or date (a = 0.05).

#### Normalized Difference Vegetation Index (NDVI)

The regressions analysis between the NDVI measured by the Field Scanalyzer and other parameters displayed a completely different pattern to the previous observation with F_v_/F_m_Sc. All the linear regressions were significant for Cadenza, whereas no significant regressions were observed for Bobwhite (Table 2; Supplementary Fig. S4). Cadenza showed high correlations for most of the parameters, regardless of the type of measurement with R ^2^ ranging from 0.55 to 0.84 for the gas-exchange parameters, 0.31 to 0.79 for the biochemical parameters, and 0.42 to 0.75 for handheld devices. The Generalist Models were clearly impacted by the lack of responsiveness of the ground variables to changes in NDVI for Bobwhite, displaying lower but still significant R ^2^_adj_ for most parameters (R^2^_adj_ = 0.16 – 0.53), except for Chl b and SPAD. However, the Interaction Models did not show a clear cultivar effect for all the parameters. Significant cv. effects were only observed for V_cmax_ among the gas-exchange parameters, for Chl a and Chl b for the biochemical parameters and for SPAD, Φ(II)FP and F_v_/F_m_HP. Incorporating the cv. effect into the Interaction Model improved the regressions in most cases (R ^2^ = 0.29 – 0.60), except for g_m_. Significant interactions were found for Chl a, b SPAD and Φ(II)FM, confirming the different behaviour of Bobwhite in its responses to NDVI (Fig. 3).

**Table 2.** Parameters of linear regressions between the normalized difference vegetation index (NDVI) measured by the Field Scanalyzer and other measured parameters.

| Parameters |  |  | Bobwhite |  |  | Cadenza |  |  | Gen. Model |  |  | Line effect |  |  |  |  |  |  |  |
| --- | --- | --- | --- | --- | --- | --- | --- | --- | --- | --- | --- | --- | --- | --- | --- | --- | --- | --- | --- |
| y |  | x | F | p | R<sup>2</sup><sub>adj</sub> | F | p | R<sup>2</sup><sub>adj</sub> | F | p | R<sup>2</sup><sub>adj</sub> | F (Par.) | p (Par.) | F (cv.) | p (cv.) | F (Int.) | p (Int.) | p (Eq) | R<sup>2</sup><sub>adj</sub> |
| V<sub>cmax</sub> | vs | NDVI | 0.7 | 0.405 | -0.02 | 43.0 | 0.000 | 0.75 | 21.2 | 0.000 | 0.41 | 25.0 | 0.000 | 4.5 | 0.043 | 2.6 | 0.121 | 0.000 | 0.50 |
| J | vs | NDVI | 0.8 | 0.379 | -0.01 | 74.0 | 0.000 | 0.84 | 31.6 | 0.000 | 0.51 | 38.2 | 0.000 | 3.8 | 0.063 | 4.1 | 0.053 | 0.000 | 0.60 |
| g<sub>m</sub> | vs | NDVI | 1.2 | 0.302 | 0.01 | 18.3 | 0.001 | 0.55 | 16.3 | 0.000 | 0.35 | 15.3 | 0.001 | 0.1 | 0.794 | 0.3 | 0.587 | 0.006 | 0.30 |
| k<sub>c</sub><sup>c</sup> | vs | NDVI | 1.2 | 0.298 | 0.01 | 36.4 | 0.000 | 0.72 | 27.3 | 0.000 | 0.48 | 28.1 | 0.000 | 1.1 | 0.307 | 1.8 | 0.188 | 0.000 | 0.49 |
| R<sub>photoresp</sub> | vs | NDVI | 0.9 | 0.368 | -0.01 | 23.2 | 0.000 | 0.61 | 24.9 | 0.000 | 0.45 | 24.7 | 0.000 | 0.0 | 0.954 | 1.7 | 0.202 | 0.000 | 0.45 |
| A<sub>max</sub> | vs | NDVI | 0.9 | 0.352 | -0.01 | 34.2 | 0.000 | 0.70 | 33.1 | 0.000 | 0.53 | 35.2 | 0.000 | 1.6 | 0.212 | 2.2 | 0.152 | 0.000 | 0.55 |
| TSP | vs | NDVI | 1.5 | 0.247 | 0.03 | 52.2 | 0.000 | 0.79 | 27.7 | 0.000 | 0.48 | 31.2 | 0.000 | 3.7 | 0.066 | 1.9 | 0.180 | 0.000 | 0.54 |
| Chl a | vs | NDVI | 3.1 | 0.101 | 0.13 | 13.3 | 0.003 | 0.47 | 6.6 | 0.016 | 0.16 | 9.5 | 0.005 | 5.0 | 0.035 | 9.4 | 0.005 | 0.001 | 0.42 |
| Chl b | vs | NDVI | 4.4 | 0.057 | 0.19 | 7.2 | 0.019 | 0.31 | 2.3 | 0.139 | 0.04 | 3.3 | 0.081 | 6.5 | 0.017 | 7.1 | 0.013 | 0.004 | 0.32 |
| Rubisco | vs | NDVI | 0.1 | 0.740 | -0.07 | 37.0 | 0.000 | 0.72 | 20.0 | 0.000 | 0.40 | 21.8 | 0.000 | 0.8 | 0.369 | 3.7 | 0.065 | 0.000 | 0.45 |
| V<sub>i</sub> | vs | NDVI | 0.4 | 0.564 | -0.05 | 46.3 | 0.000 | 0.76 | 26.3 | 0.000 | 0.47 | 28.4 | 0.000 | 0.7 | 0.426 | 3.6 | 0.069 | 0.000 | 0.51 |
| V<sub>t</sub> | vs | NDVI | 0.6 | 0.443 | -0.03 | 49.2 | 0.000 | 0.78 | 22.0 | 0.000 | 0.42 | 25.5 | 0.000 | 3.8 | 0.062 | 2.7 | 0.110 | 0.000 | 0.50 |
| SPAD | vs | NDVI | 0.6 | 0.467 | -0.03 | 14.6 | 0.002 | 0.49 | 3.9 | 0.059 | 0.09 | 7.1 | 0.013 | 18.7 | 0.000 | 6.4 | 0.017 | 0.000 | 0.50 |
| Φ(II)FP | vs | NDVI | 0.4 | 0.545 | -0.05 | 11.2 | 0.005 | 0.42 | 18.9 | 0.000 | 0.38 | 22.0 | 0.000 | 4.6 | 0.042 | 2.0 | 0.170 | 0.000 | 0.47 |
| Φ(II)FM | vs | NDVI | 0.1 | 0.808 | -0.07 | 29.9 | 0.000 | 0.67 | 14.3 | 0.001 | 0.32 | 18.7 | 0.000 | 3.6 | 0.069 | 6.8 | 0.015 | 0.000 | 0.47 |
| Φ(II)HP | vs | NDVI | 0.0 | 0.927 | -0.08 | 13.0 | 0.003 | 0.46 | 9.6 | 0.004 | 0.23 | 10.5 | 0.003 | 3.2 | 0.084 | 1.3 | 0.261 | 0.007 | 0.29 |
| F<sub>v</sub>/F<sub>m</sub>HP | vs | NDVI | 0.6 | 0.466 | -0.03 | 41.9 | 0.000 | 0.75 | 29.9 | 0.000 | 0.50 | 35.5 | 0.000 | 4.9 | 0.036 | 2.3 | 0.138 | 0.000 | 0.58 |

y and x, elements of the regression; p (Eq), p-value of the regression; R<sup>2</sup><sub>adj</sub>, adjusted coefficient of determination of the regression; p (Eq) < 0.05, the regression is significant (rows in black);

### Phenocenter

#### Dark-adapted chlorophyll fluorescence (F_v_/F_m_Pc)

The F_v_/F_m_Pc calculated through the IMAGING-PAM at the Phenocenter for the flag leaves (Fig. 1) also showed that the linear regressions were strongly affected by the cultivar (Table 3; Supplementary Fig. S5). Cadenza demonstrated high and significant correlations with all the parameters measured by gas exchange (R ^2^_adj_ = 0.57 – 0.85), biochemical analysis (R ^2^_adj_ = 0.35 – 0.76) or handheld device (R^2^_adj_ = 0.50 ‒ 0.82), with the highest fits observed for J, Φ(II)FM and F_v_/F_m_HP (R^2^_adj_ = 0.85, 0.81 and 0.82, respectively) and the lowest fits for Chl a and b (R ^2^_adj_ = 0.44 and 0.35, respectively). For Bobwhite, significant regressions were only observed between F_v_/F_m_Pc and the parameters from the biochemical analysis, Chl a and Chl b and the parameters from the handheld devices, Φ(II)HP (R ^2^_adj_ = 0.32, 0.25 and 0.45, respectively). All the regression showed a significant R ^2^_adj_ for the Generalist Models. Since most of the regressions for Bobwhite were non-significant, it was expected that a cultivar effect would be present for most parameters.However, the Interaction Models only showed a significant cv. effect for A_max_, SPAD, Φ(II)FP, Φ(II)HP and F_v_/F_m_HP. This indicates that for these five parameters, the offset of the regression differs significantly between cultivars. The only significant interaction was found for Φ(II)HP, indicating that in this case, the slopes were significantly different between cultivars only for Φ(II)HP. In general, the R ^2^_adj_ between the Generalist and the Interaction Models were similar, with the exception of the five parameters that had a significant cv. effect. For those, improvements to the regressions were achieved by considering the cv. effect into the Interaction Models.

**Table 3.**
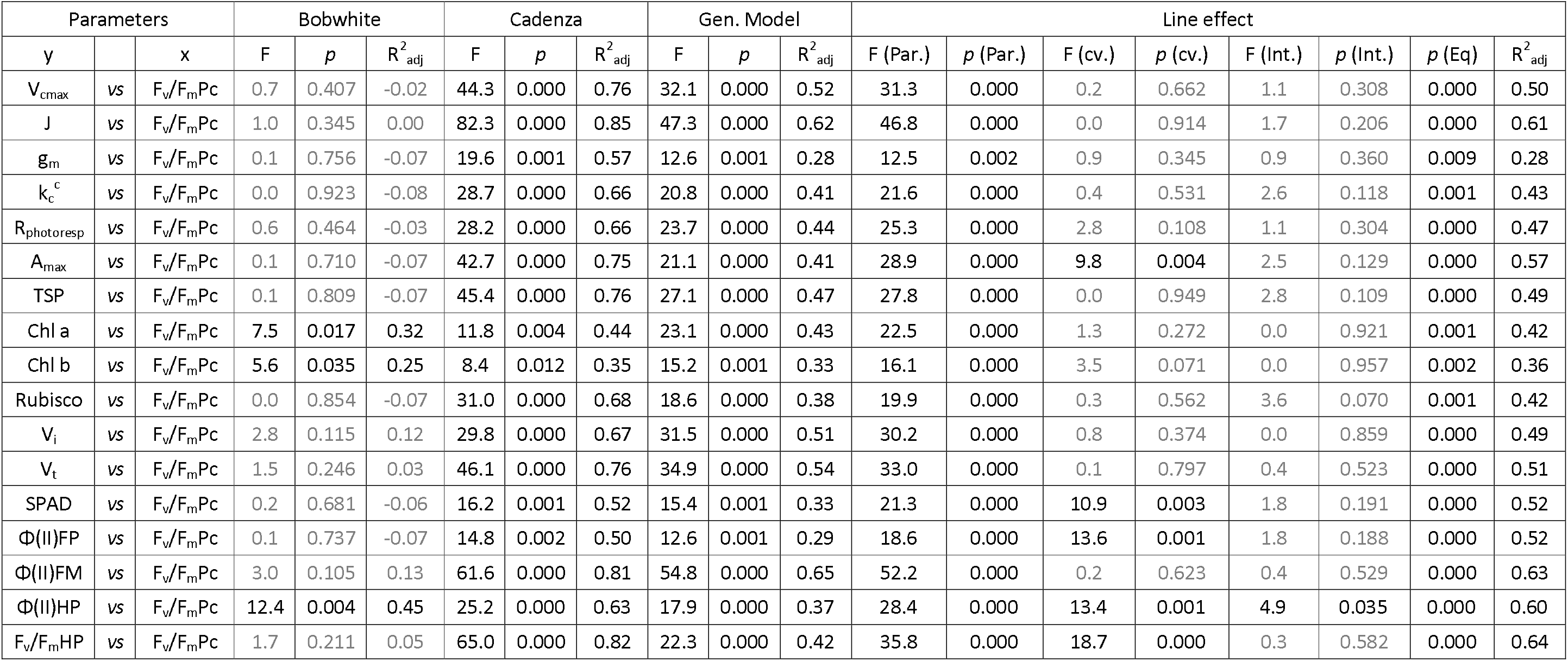
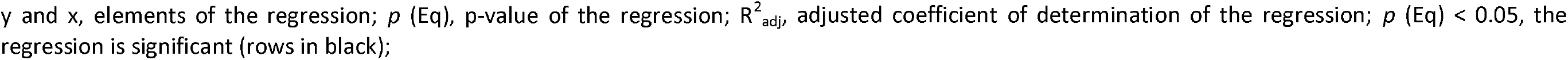
Parameters of linear regressions between chlorophyll fluorescence (F_v_/F_m_) measured by the Phenocenter and other measured parameters.

| Parameters |  |  | Bobwhite |  |  | Cadenza |  |  | Gen. Model |  |  | Line effect |  |  |  |  |  |  |  |
| --- | --- | --- | --- | --- | --- | --- | --- | --- | --- | --- | --- | --- | --- | --- | --- | --- | --- | --- | --- |
| y |  | x | F | p | R<sup>2</sup><sub>adj</sub> | F | p | R<sup>2</sup><sub>adj</sub> | F | p | R<sup>2</sup><sub>adj</sub> | F (Par.) | p (Par.) | F (cv.) | p (cv.) | F (Int.) | p (Int.) | p (Eq) | R<sup>2</sup><sub>adj</sub> |
| V<sub>cmax</sub> | vs | $F_v/F_mPc$ | 0.7 | 0.407 | -0.02 | 44.3 | 0.000 | 0.76 | 32.1 | 0.000 | 0.52 | 31.3 | 0.000 | 0.2 | 0.662 | 1.1 | 0.308 | 0.000 | 0.50 |
| J | vs | $F_v/F_mPc$ | 1.0 | 0.345 | 0.00 | 82.3 | 0.000 | 0.85 | 47.3 | 0.000 | 0.62 | 46.8 | 0.000 | 0.0 | 0.914 | 1.7 | 0.206 | 0.000 | 0.61 |
| g<sub>m</sub> | vs | $F_v/F_mPc$ | 0.1 | 0.756 | -0.07 | 19.6 | 0.001 | 0.57 | 12.6 | 0.001 | 0.28 | 12.5 | 0.002 | 0.9 | 0.345 | 0.9 | 0.360 | 0.009 | 0.28 |
| k<sub>c</sub><sup>c</sup> | vs | $F_v/F_mPc$ | 0.0 | 0.923 | -0.08 | 28.7 | 0.000 | 0.66 | 20.8 | 0.000 | 0.41 | 21.6 | 0.000 | 0.4 | 0.531 | 2.6 | 0.118 | 0.001 | 0.43 |
| R<sub>photoresp</sub> | vs | $F_v/F_mPc$ | 0.6 | 0.464 | -0.03 | 28.2 | 0.000 | 0.66 | 23.7 | 0.000 | 0.44 | 25.3 | 0.000 | 2.8 | 0.108 | 1.1 | 0.304 | 0.000 | 0.47 |
| A<sub>max</sub> | vs | $F_v/F_mPc$ | 0.1 | 0.710 | -0.07 | 42.7 | 0.000 | 0.75 | 21.1 | 0.000 | 0.41 | 28.9 | 0.000 | 9.8 | 0.004 | 2.5 | 0.129 | 0.000 | 0.57 |
| TSP | vs | $F_v/F_mPc$ | 0.1 | 0.809 | -0.07 | 45.4 | 0.000 | 0.76 | 27.1 | 0.000 | 0.47 | 27.8 | 0.000 | 0.0 | 0.949 | 2.8 | 0.109 | 0.000 | 0.49 |
| Chl a | vs | $F_v/F_mPc$ | 7.5 | 0.017 | 0.32 | 11.8 | 0.004 | 0.44 | 23.1 | 0.000 | 0.43 | 22.5 | 0.000 | 1.3 | 0.272 | 0.0 | 0.921 | 0.001 | 0.42 |
| Chl b | vs | $F_v/F_mPc$ | 5.6 | 0.035 | 0.25 | 8.4 | 0.012 | 0.35 | 15.2 | 0.001 | 0.33 | 16.1 | 0.000 | 3.5 | 0.071 | 0.0 | 0.957 | 0.002 | 0.36 |
| Rubisco | vs | $F_v/F_mPc$ | 0.0 | 0.854 | -0.07 | 31.0 | 0.000 | 0.68 | 18.6 | 0.000 | 0.38 | 19.9 | 0.000 | 0.3 | 0.562 | 3.6 | 0.070 | 0.001 | 0.42 |
| V<sub>i</sub> | vs | $F_v/F_mPc$ | 2.8 | 0.115 | 0.12 | 29.8 | 0.000 | 0.67 | 31.5 | 0.000 | 0.51 | 30.2 | 0.000 | 0.8 | 0.374 | 0.0 | 0.859 | 0.000 | 0.49 |
| V<sub>t</sub> | vs | $F_v/F_mPc$ | 1.5 | 0.246 | 0.03 | 46.1 | 0.000 | 0.76 | 34.9 | 0.000 | 0.54 | 33.0 | 0.000 | 0.1 | 0.797 | 0.4 | 0.523 | 0.000 | 0.51 |
| SPAD | vs | $F_v/F_mPc$ | 0.2 | 0.681 | -0.06 | 16.2 | 0.001 | 0.52 | 15.4 | 0.001 | 0.33 | 21.3 | 0.000 | 10.9 | 0.003 | 1.8 | 0.191 | 0.000 | 0.52 |
| Φ(II)FP | vs | $F_v/F_mPc$ | 0.1 | 0.737 | -0.07 | 14.8 | 0.002 | 0.50 | 12.6 | 0.001 | 0.29 | 18.6 | 0.000 | 13.6 | 0.001 | 1.8 | 0.188 | 0.000 | 0.52 |
| Φ(II)FM | vs | $F_v/F_mPc$ | 3.0 | 0.105 | 0.13 | 61.6 | 0.000 | 0.81 | 54.8 | 0.000 | 0.65 | 52.2 | 0.000 | 0.2 | 0.623 | 0.4 | 0.529 | 0.000 | 0.63 |
| Φ(II)HP | vs | $F_v/F_mPc$ | 12.4 | 0.004 | 0.45 | 25.2 | 0.000 | 0.63 | 17.9 | 0.000 | 0.37 | 28.4 | 0.000 | 13.4 | 0.001 | 4.9 | 0.035 | 0.000 | 0.60 |
| $F_v/F_mHP$ | vs | $F_v/F_mPc$ | 1.7 | 0.211 | 0.05 | 65.0 | 0.000 | 0.82 | 22.3 | 0.000 | 0.42 | 35.8 | 0.000 | 18.7 | 0.000 | 0.3 | 0.582 | 0.000 | 0.64 |

y and x, elements of the regression; $p$ (Eq), p-value of the regression; $R^2_{adj}$ , adjusted coefficient of determination of the regression; $p$ (Eq) < 0.05, the regression is significant (rows in black);

#### Photosystem II quantum efficiency (Φ(II)_532_)

The Φ(II)_532_ was calculated through the Phenocenter IMAGING-PAM, at 532 µmols m ^-2^ s ^-1^ photons during a light curve at flag leaf level (Fig. 1). The regressions between Φ(II)_532_ and most of the parameters displayed a significant cultivar effect except for Chl a, b and SPAD (Table 4, Supplementary Fig. S6). Cadenza presented significant correlations with all the parameters measured either by gas-exchange (R ^2^_adj_ = 0.59 – 0.84), biochemical analysis (R ^2^_adj_ = 0.32 – 0.83) or handheld device (R^2^_adj_ = 0.31 – 0.90), with the highest fittings for J, A_max_, TSP, V_t_ and F_v_/F_m_HP (R^2^_adj_ = 0.82, 0.84, 0.82, 0.83 and 0.90, respectively) and the lowest fittings for Chl b and Φ(II)FM (R ^2^_adj_ = 0.32 and 0.31, respectively). For Bobwhite, the regressions between Φ(II)_532_ and the gas-exchange parameters were all significant (R ^2^_adj_ = 0.36 – 0.63), whereas the regression with the biochemical parameters and the handheld devices parameters were only significant for TSP, V_i_, V_t_, Φ(II)FP and F_v_/F_m_HP (R ^2^_adj_ = 0.37, 0.27, 0.31, 0.28 and 0.34, respectively). As already indicated, most of the parameters (except for Chl a, b, and SPAD) showed a significant cv. effect, indicating that the intercept of linear regressions for each of the cultivars was significantly different. Among those parameters V_cmax_, g_m_, k_c_^c^, R_photoresp_, TSP, Rubisco, V_i_, Φ(II)FP and Φ(II)HP did not show significant interactions, indicating that the slopes of the linear regression for each cultivar were not significantly different. The relationship between Φ(II)_532_ and those nine parameters followed a similar pattern, with only intercepts differing between cultivar. As expected, the Generalist Models were affected by the cultivar. The R ^2^_adj_ was not significant for A_max_, Φ(II)FP, Φ(II)HP and F_v_/F_m_HP, and was low to moderate, but significant, for the other parameters (R ^2^_adj_ = 0.14 – 0.44). Considering the cultivar effect in the Interaction Model substantially improved the models for all the parameters, with R ^2^_adj_ ranging from 0.32 to 0.77.

**Table 4.** Parameters of linear regressions between photosystem II quantum efficiency (Φ(II)_532_) measured by the Phenocenter and other measured parameters.

| Parameters |  |  | Bobwhite |  |  | Cadenza |  |  | Gen. Model |  |  | Line effect |  |  |  |  |  |  |  |
| --- | --- | --- | --- | --- | --- | --- | --- | --- | --- | --- | --- | --- | --- | --- | --- | --- | --- | --- | --- |
| y | | x | F | $p$ | $R^2_{adj}$ | F | $p$ | $R^2_{adj}$ | F | $p$ | $R^2_{adj}$ | F (Par.) | $p$ (Par.) | F (cv.) | $p$ (cv.) | F (Int.) | $p$ (Int.) | $p$ (Eq) | $R^2_{adj}$ |
| $V_{cmax}$ | vs | $\Phi(II)_{532}$ | 19.2 | 0.001 | 0.57 | 34.0 | 0.000 | 0.70 | 23.5 | 0.000 | 0.44 | 37.8 | 0.000 | 17.6 | 0.000 | 1.4 | 0.244 | 0.000 | 0.65 |
| J | vs | $\Phi(II)_{532}$ | 17.9 | 0.001 | 0.55 | 65.9 | 0.000 | 0.82 | 21.6 | 0.000 | 0.42 | 48.5 | 0.000 | 32.3 | 0.000 | 4.6 | 0.041 | 0.000 | 0.74 |
| $g_m$ | vs | $\Phi(II)_{532}$ | 15.4 | 0.002 | 0.51 | 26.6 | 0.000 | 0.65 | 9.9 | 0.004 | 0.23 | 18.0 | 0.000 | 24.9 | 0.000 | 0.0 | 0.850 | 0.000 | 0.58 |
| $k_c^c$ | vs | $\Phi(II)_{532}$ | 24.9 | 0.000 | 0.63 | 21.0 | 0.001 | 0.59 | 12.7 | 0.001 | 0.29 | 22.0 | 0.000 | 22.0 | 0.000 | 0.5 | 0.478 | 0.000 | 0.59 |
| $R_{photoresp}$ | vs | $\Phi(II)_{532}$ | 16.4 | 0.001 | 0.52 | 41.2 | 0.000 | 0.74 | 8.6 | 0.007 | 0.21 | 21.7 | 0.000 | 41.5 | 0.000 | 2.8 | 0.108 | 0.000 | 0.68 |
| $A_{max}$ | vs | $\Phi(II)_{532}$ | 9.0 | 0.010 | 0.36 | 73.2 | 0.000 | 0.84 | 3.9 | 0.057 | 0.09 | 13.4 | 0.001 | 64.0 | 0.000 | 5.5 | 0.027 | 0.000 | 0.73 |
| TSP | vs | $\Phi(II)_{532}$ | 9.3 | 0.009 | 0.37 | 65.0 | 0.000 | 0.82 | 19.6 | 0.000 | 0.39 | 36.3 | 0.000 | 21.9 | 0.000 | 4.1 | 0.052 | 0.000 | 0.67 |
| Chl a | vs | $\Phi(II)_{532}$ | 0.9 | 0.370 | -0.01 | 13.0 | 0.003 | 0.46 | 7.0 | 0.013 | 0.17 | 9.5 | 0.005 | 1.0 | 0.328 | 11.4 | 0.002 | 0.001 | 0.39 |
| Chl b | vs | $\Phi(II)_{532}$ | 3.2 | 0.095 | 0.14 | 7.5 | 0.017 | 0.32 | 5.7 | 0.024 | 0.14 | 7.2 | 0.012 | 0.1 | 0.802 | 9.6 | 0.005 | 0.004 | 0.32 |
| Rubisco | vs | $\Phi(II)_{532}$ | 4.2 | 0.061 | 0.19 | 30.9 | 0.000 | 0.68 | 9.4 | 0.005 | 0.22 | 14.7 | 0.001 | 14.8 | 0.001 | 3.0 | 0.094 | 0.000 | 0.50 |
| $V_i$ | vs | $\Phi(II)_{532}$ | 6.3 | 0.026 | 0.27 | 43.1 | 0.000 | 0.75 | 10.3 | 0.003 | 0.24 | 19.3 | 0.000 | 22.7 | 0.000 | 3.8 | 0.062 | 0.000 | 0.60 |
| $V_t$ | vs | $\Phi(II)_{532}$ | 7.3 | 0.018 | 0.31 | 69.7 | 0.000 | 0.83 | 20.1 | 0.000 | 0.40 | 34.8 | 0.000 | 18.3 | 0.000 | 4.3 | 0.048 | 0.000 | 0.65 |
| SPAD | vs | $\Phi(II)_{532}$ | 0.3 | 0.609 | -0.05 | 21.6 | 0.000 | 0.60 | 18.9 | 0.000 | 0.38 | 27.2 | 0.000 | 0.1 | 0.713 | 14.2 | 0.001 | 0.000 | 0.57 |
| $\Phi(II)_{FP}$ | vs | $\Phi(II)_{532}$ | 6.5 | 0.024 | 0.28 | 7.2 | 0.019 | 0.31 | 0.2 | 0.623 | -0.03 | 0.4 | 0.510 | 23.6 | 0.000 | 1.2 | 0.281 | 0.000 | 0.43 |
| $\Phi(II)_{FM}$ | vs | $\Phi(II)_{532}$ | 2.6 | 0.132 | 0.10 | 47.0 | 0.000 | 0.77 | 15.6 | 0.000 | 0.34 | 25.4 | 0.000 | 11.3 | 0.002 | 8.2 | 0.008 | 0.000 | 0.59 |
| $\Phi(II)_{HP}$ | vs | $\Phi(II)_{532}$ | 0.9 | 0.371 | -0.01 | 25.5 | 0.000 | 0.64 | 0.4 | 0.545 | -0.02 | 0.6 | 0.435 | 19.2 | 0.000 | 1.9 | 0.184 | 0.001 | 0.39 |
| $F_v/F_{mHP}$ | vs | $\Phi(II)_{532}$ | 8.3 | 0.013 | 0.34 | 131.7 | 0.000 | 0.90 | 2.2 | 0.153 | 0.04 | 8.9 | 0.006 | 84.4 | 0.000 | 4.7 | 0.040 | 0.000 | 0.77 |

y and x, elements of the regression; $p$ (Eq), p-value of the regression; $R^2_{adj}$ , adjusted coefficient of determination of the regression; $p$ (Eq) < 0.05, the regression is significant (rows in black);

#### Non-photochemical quenching (Φ(NPQ)_1077_)

The Φ(NPQ)_1077_, calculated using the Phenocenter IMAGING-PAM, at 1077 µmols m ^-2^ s ^-1^ photons during a light curve at flag leaf level was showing significant linear regressions for most of the parameters and for both cultivars (Table 5; Supplementary Fig. S7). However, in this case, Cadenza presented significant correlations for only a few parameters measured (i) by gas exchange: V_cmax_, J, R_photoresp_ and A_max_ (R^2^_adj_ = 0.24 – 0.49), (ii) by biochemical analysis: TSP, Rubisco, V_i_ and V_t_ (R^2^_adj_ = 0.35 ‒ 0.46), and by handheld device: SPAD, Φ(II)FM, Φ(II)HP and F_v_/F_m_HP (R^2^_adj_ = 0.28 – 0.54) with the best fits for J, A_max_, TSP, SPAD and F_v_/F_m_HP (R^2^_adj_ = 0.49, 0.47, 0.46, 0.47 and 0.54, respectively). For Bobwhite, the regressions with (i) the gas-exchange parameters were all significant (R ^2^_adj_ = 0.78 - 0.89), (ii) the biochemical parameters were mostly significant (R ^2^_adj_ = 0.36 – 0.76), except for the Chl a and b content, and (iii) the handheld devices were significant only for Φ(II)FP, Φ(II)FM and F_v_/F_m_HP (R^2^_adj_ = 0.25, 0.21 and 0.65, respectively). It should be noted that for Φ(NPQ)_1077_, the R ^2^ were higher for Bobwhite for the gas-exchange and biochemical parameters, as well as for F_v_/F_m_HP of the handheld devices. The Generalist Models showed signification R ^2^_adj_ for most of the parameters (R ^2^_adj_ = 0.23 – 0.63), except for Chl a, b and SPAD. The Interaction Models showed significant cv. effects for V_cmax_, J, TSP, Chl a and b, V_t_, SPAD and Φ(II)FM, indicating a significantly difference between the two cultivars. However, for these parameters, except for SPAD, the interactions were not significant, indicating a similar slope for the regressions for both cultivars. Nonetheless, including the cv. effect in the Interaction Models improved the R ^2^_adj_ for the aforementioned parameters compared to the Generalist Models, whilst having little effect on the R ^2^_adj_ for the other parameters compared to the Generalist Models.

**Table 5.**
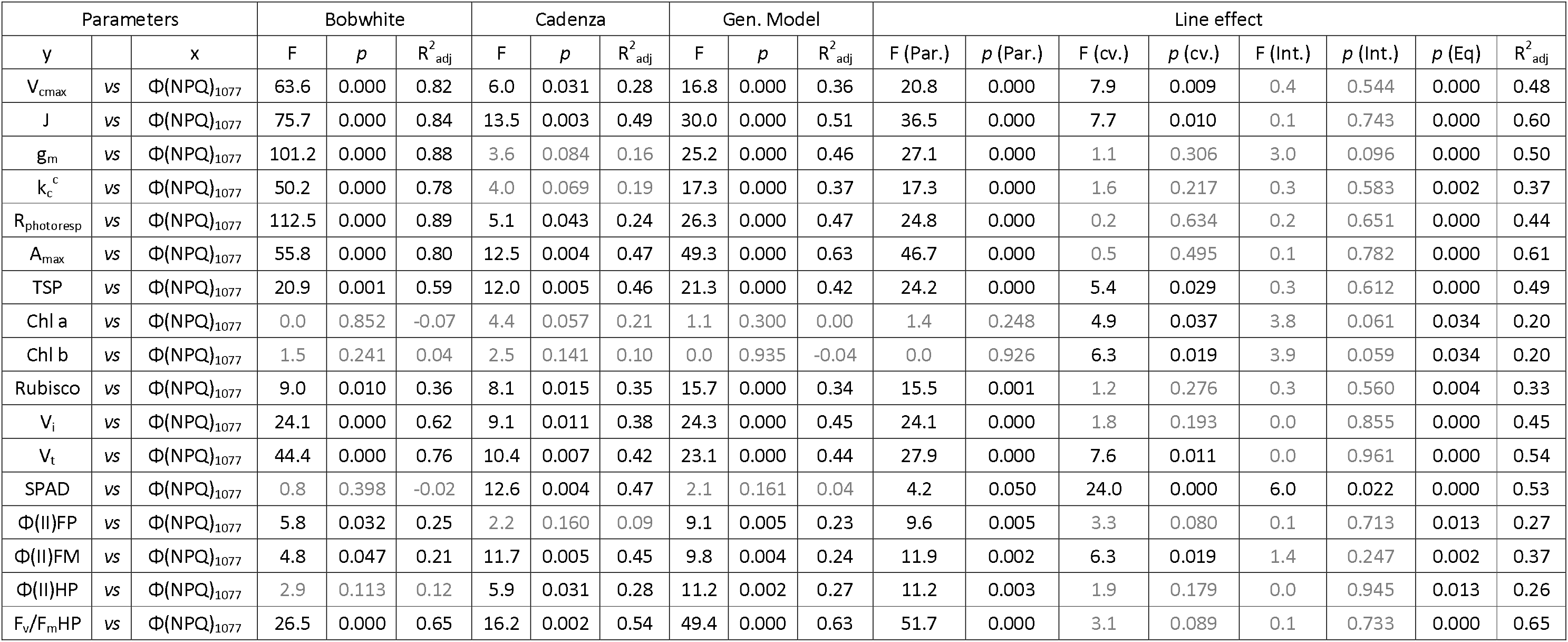
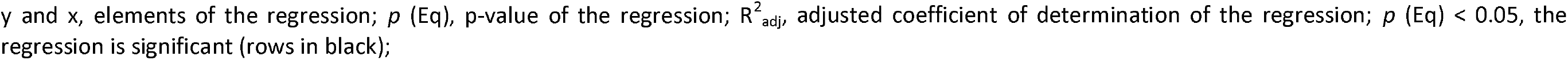
Parameters of linear regressions between non-photochemical quenching (Φ(NPQ)_1077_) measured by the Phenocenter and other measured parameters.

| Parameters |  |  | Bobwhite |  |  | Cadenza |  |  | Gen. Model |  |  | Line effect |  |  |  |  |  |  |  |
| --- | --- | --- | --- | --- | --- | --- | --- | --- | --- | --- | --- | --- | --- | --- | --- | --- | --- | --- | --- |
| y | | x | F | p | $R^2_{adj}$ | F | p | $R^2_{adj}$ | F | p | $R^2_{adj}$ | F (Par.) | p (Par.) | F (cv.) | p (cv.) | F (Int.) | p (Int.) | p (Eq) | $R^2_{adj}$ |
| $V_{cmax}$ | vs | $\Phi(NPQ)_{1077}$ | 63.6 | 0.000 | 0.82 | 6.0 | 0.031 | 0.28 | 16.8 | 0.000 | 0.36 | 20.8 | 0.000 | 7.9 | 0.009 | 0.4 | 0.544 | 0.000 | 0.48 |
| J | vs | $\Phi(NPQ)_{1077}$ | 75.7 | 0.000 | 0.84 | 13.5 | 0.003 | 0.49 | 30.0 | 0.000 | 0.51 | 36.5 | 0.000 | 7.7 | 0.010 | 0.1 | 0.743 | 0.000 | 0.60 |
| $g_m$ | vs | $\Phi(NPQ)_{1077}$ | 101.2 | 0.000 | 0.88 | 3.6 | 0.084 | 0.16 | 25.2 | 0.000 | 0.46 | 27.1 | 0.000 | 1.1 | 0.306 | 3.0 | 0.096 | 0.000 | 0.50 |
| $k_c^c$ | vs | $\Phi(NPQ)_{1077}$ | 50.2 | 0.000 | 0.78 | 4.0 | 0.069 | 0.19 | 17.3 | 0.000 | 0.37 | 17.3 | 0.000 | 1.6 | 0.217 | 0.3 | 0.583 | 0.002 | 0.37 |
| $R_{photoresp}$ | vs | $\Phi(NPQ)_{1077}$ | 112.5 | 0.000 | 0.89 | 5.1 | 0.043 | 0.24 | 26.3 | 0.000 | 0.47 | 24.8 | 0.000 | 0.2 | 0.634 | 0.2 | 0.651 | 0.000 | 0.44 |
| $A_{max}$ | vs | $\Phi(NPQ)_{1077}$ | 55.8 | 0.000 | 0.80 | 12.5 | 0.004 | 0.47 | 49.3 | 0.000 | 0.63 | 46.7 | 0.000 | 0.5 | 0.495 | 0.1 | 0.782 | 0.000 | 0.61 |
| TSP | vs | $\Phi(NPQ)_{1077}$ | 20.9 | 0.001 | 0.59 | 12.0 | 0.005 | 0.46 | 21.3 | 0.000 | 0.42 | 24.2 | 0.000 | 5.4 | 0.029 | 0.3 | 0.612 | 0.000 | 0.49 |
| Chl a | vs | $\Phi(NPQ)_{1077}$ | 0.0 | 0.852 | -0.07 | 4.4 | 0.057 | 0.21 | 1.1 | 0.300 | 0.00 | 1.4 | 0.248 | 4.9 | 0.037 | 3.8 | 0.061 | 0.034 | 0.20 |
| Chl b | vs | $\Phi(NPQ)_{1077}$ | 1.5 | 0.241 | 0.04 | 2.5 | 0.141 | 0.10 | 0.0 | 0.935 | -0.04 | 0.0 | 0.926 | 6.3 | 0.019 | 3.9 | 0.059 | 0.034 | 0.20 |
| Rubisco | vs | $\Phi(NPQ)_{1077}$ | 9.0 | 0.010 | 0.36 | 8.1 | 0.015 | 0.35 | 15.7 | 0.000 | 0.34 | 15.5 | 0.001 | 1.2 | 0.276 | 0.3 | 0.560 | 0.004 | 0.33 |
| $V_i$ | vs | $\Phi(NPQ)_{1077}$ | 24.1 | 0.000 | 0.62 | 9.1 | 0.011 | 0.38 | 24.3 | 0.000 | 0.45 | 24.1 | 0.000 | 1.8 | 0.193 | 0.0 | 0.855 | 0.000 | 0.45 |
| $V_t$ | vs | $\Phi(NPQ)_{1077}$ | 44.4 | 0.000 | 0.76 | 10.4 | 0.007 | 0.42 | 23.1 | 0.000 | 0.44 | 27.9 | 0.000 | 7.6 | 0.011 | 0.0 | 0.961 | 0.000 | 0.54 |
| SPAD | vs | $\Phi(NPQ)_{1077}$ | 0.8 | 0.398 | -0.02 | 12.6 | 0.004 | 0.47 | 2.1 | 0.161 | 0.04 | 4.2 | 0.050 | 24.0 | 0.000 | 6.0 | 0.022 | 0.000 | 0.53 |
| $\Phi(II)FP$ | vs | $\Phi(NPQ)_{1077}$ | 5.8 | 0.032 | 0.25 | 2.2 | 0.160 | 0.09 | 9.1 | 0.005 | 0.23 | 9.6 | 0.005 | 3.3 | 0.080 | 0.1 | 0.713 | 0.013 | 0.27 |
| $\Phi(II)FM$ | vs | $\Phi(NPQ)_{1077}$ | 4.8 | 0.047 | 0.21 | 11.7 | 0.005 | 0.45 | 9.8 | 0.004 | 0.24 | 11.9 | 0.002 | 6.3 | 0.019 | 1.4 | 0.247 | 0.002 | 0.37 |
| $\Phi(II)HP$ | vs | $\Phi(NPQ)_{1077}$ | 2.9 | 0.113 | 0.12 | 5.9 | 0.031 | 0.28 | 11.2 | 0.002 | 0.27 | 11.2 | 0.003 | 1.9 | 0.179 | 0.0 | 0.945 | 0.013 | 0.26 |
| $F_v/F_{mHP}$ | vs | $\Phi(NPQ)_{1077}$ | 26.5 | 0.000 | 0.65 | 16.2 | 0.002 | 0.54 | 49.4 | 0.000 | 0.63 | 51.7 | 0.000 | 3.1 | 0.089 | 0.1 | 0.733 | 0.000 | 0.65 |

y and x, elements of the regression; $p$ (Eq), p-value of the regression; $R^2_{adj}$ , adjusted coefficient of determination of the regression; $p$ (Eq) < 0.05, the regression is significant (rows in black);

## DISCUSSION

### Model performances and cultivar effect

The present study aimed to model photosynthetic parameters collected at leaf scale through gas-exchange and by biochemical analysis using imaging technology. Five image variables collected at pot scale in the field (NDVI and F_v_/F_m_Sc) and at leaf scale in laboratory conditions (F_v_/F_m_Pc, Φ(II)_532_ and Φ(NPQ)_1077_) were investigated as potential targets for further screening of photosynthetic parameters in phenotyping trials. Overall, the results showed the possibilities to build linear models to predict photosynthesis parameters based on the image variables, despite the varying performance of the two cultivars. The Interaction Models proved that considering the cultivar effect improved the models R ^2^_adj_ when the cultivar effect was significant compared to the Generalist Models. The linear regression model performances were largely driven by the phenotypic variability resulting from the stage of development of the two cultivars. As senescence progressed, a decrease of the phenotypic values for each cultivar was observed for all the gas exchange parameters (V_cmax_, J, g_m_, k_c_^c^, R_photoresp_ and A_max_) as well as the Rubisco related parameters (TSP, Rubisco, V_i_, V_t_), the fluorescence parameters F_v_/F_m_HP collected with the HandiPEA and Φ(NPQ)_1077_ measured at the Phenocenter. A contrasting behaviour was observed between Bobwhite and Cadenza for chlorophyll variables (Chl a and b, SPAD) and the Φ(II) parameters measured by the handheld devices. The same was observed for the other four image variables. Cadenza followed the same senescence pattern with a decrease in phenotypic values with the senescence, explaining the good agreement between the image and the ground variables. The lower model performances with Φ(NPQ)_1077_ were due to the fact that a significant decrease of NPQ could only be seen at 21 days post anthesis (DPA). The low model performances between the five image variables and the chlorophyll index resulted from the fact that the chlorophyll amount increased up to 7 DPA before decreasing. In contrast, Bobwhite displayed a different pattern with no significant variation of the phenotypic values across the five dates of measurement for the chlorophyll variables (Chl a and b, SPAD), the Φ(II) variables as well as for F_v_/F_m_Sc, NDVI, F_v_/F_m_Pc. While no significant decreases were observed for F_v_/F_m_Sc, a decreasing trend was still observed, which explains why the model performances were similar to Cadenza. A decreasing trend was also observed for NDVI without significant differences between dates. However, the small day-to-day fluctuations explained the absence of correlation with the photosynthetic variables. The absence of trend and significant differences between dates of F_v_/F_m_Pc explained the absence of correlations with the ground variables for Bobwhite. The relatively good performance of Bobwhite for Φ(II)_532_ seems to be due to the first date having a significant higher value than the four other dates.

The observed patterns in Cadenza correspond to a typical wheat senescence pattern with a gradual decrease in photosynthetic capacity along with the degradation of the chlorophyll (Thomas & Howarth, 2000). On the other hand, Bobwhite is known as a slow maturaturing cultivar (Mackintosh *et al*., 2006), and it was expected to see delayed decrease in phenotypic values for both photosynthesis and chlorophyll variables, or a slower maturation compared to Cadenza, which would correspond to a Type A or B stay green phenotype according to Thomas & Howarth (2000). However, the current results suggest Bobwhite may belong to Type C (also called cosmetic stay-green) which is described as undergoing a degradation of the photosynthetic activity as normal with, however, no sign of degradation of the chlorophyll, which corresponds to the results observed in this study, at least up to the end of the experiment at 21 days after anthesis. According to Thomas and Ougham (2014), breeders tend to avoid using cosmetic stay-greens because they do not offer any yield benefits. The opposite type, functional stay-green, is more desirable for breeding purposes as photosynthesis is maintained longer during the maturation phase (Thomas & Ougham, 2014; Li *et al*., 2022). However, screening populations for functional stray-green genotypes is complicated as it would require gas-exchange measurements at multiple time points on a large number of individuals, which is not feasible without a large number of instruments and trained users. Currently, phenotyping studies on stay-green genotypes are based on screening materials with visual scoring, SPAD, RGB images or spectral indices, without being able to distinguish cosmetic from functional genotypes (Pinto *et al*., 2016; Christopher *et al*., 2018, Hassan *et al*., 2021, Chapman *et al*., 2021). The last five years have seen successful studies predicting V_cmax_ and J using spectral data (Silva-Perez *et al*., 2018; Meacham-Hensold *et al*., 2019, Montes *et al*., 2022). It would be interesting to see if those approaches can overcome the challenges of identifying functional stay green phenotype in field conditions.

### Image variables vs traditional approaches

The results showed the image variables, tested at the plant scale (NDVI and F_v_/F_m_Sc), and at the leaf scale (F_v_/F_m_Pc, Φ(II)_532_ and Φ(NPQ)_1077_) constitute good to strong estimators of the gas-exchange (V_cmax_, J, g_m_, k_c_^c^, R_photoresp_ and A_max_) and the biochemical variables (TSP, Rubisco, V_i_ and V_t_), when senescence is driving the variability. As mentioned above, the model performances have been strongly influenced by the cultivar disparity in term of the senescence patterns, resulting in strong specificity between the ground variables and their relationship with the image-based variables. The highest predictions for the gas-exchange variables were obtained with Φ(NPQ)_1077_ (R^2^_adj_ = 0.78-0.89) for the cultivar Bobwhite. NDVI, F_v_/F_m_Pc and Φ(II)_532_ were also strong estimators for V_cmax_, J and A_max_ with the cultivar Cadenza. Lower performances were obtained using F_v_/F_m_Sc but with acceptable results (R^2^_adj_ ∼ 0.50). However, it was not possible to conclude which of the image variables was the most relevant to investigate a particular parameter in the photosynthetic measurements collected by gas-exchange or biochemical analysis.

NDVI has been used extensively over decades to monitor vegetation dynamics and quantify green biomass at various scales (landscape to plot). It is well accepted that NDVI is not the most suitable index to monitor instantaneous change in net CO_2_ assimilation (Gamon *et al*., 1995, Dobrowski *et al*., 2005; Urban *et al*., 2013). However, a few studies showed significant relationship between NDVI and net CO_2_ assimilation, mostly driven by the genetic variability as well as by the stages of development, similarly to our study (Kumari *et al*., 2012; Gonzalez-Dugo *et al*., 2015; Camino *et al*., 2018). Studies in the literature looking at the link between NDVI and gas-exchange and biochemical parameters are scarce to our knowledge. Letts *et al*. (2008) found an absence of correlation between A_max_ and NDVI while Zhou *et al*. (2014) showed a high correlation with V_cmax_ on deciduous forest trees, mainly due to the different phenological stages. Despite the absence of correlation observed for Bobwhite, due to the reasons discussed, the Interaction Models showed that it was possible to use NDVI to predict the different parameters related to carbon assimilation, either derived from A-C_i_ and A-Par curve (V_cmax_, J, g_m_, k_c_^c^, R_photoresp_, A_max_) or from biochemical analysis (V_i_ and V_t_) but not the Rubisco activation state (Act, Table S3). The same was observed for N related parameters (TSP and Rubisco) and chlorophyll pigment (Chl a and b, SPAD). Despite the strong discrepancy existing between the two cultivars caused by their senescence behaviour, the Interaction Models showed that NDVI could be a predictor for those variables with acceptable results. Gitelson *et al*. (1998, 2003) showed that NDVI was far from the best index to predict chlorophyll content at leaf scale due to its lack of sensitivity from medium chlorophyll concentration. However, other studies proved that, at the canopy scale, NDVI might be a useful indicator if no other indices are available (Zarco-Tejada *et al*., 2005; Camino *et al*., 2018; Alordzinu *et al*., 2021).

The maximum efficiency of the photosystem II (F_v_/F_m_) is commonly used as an indicator of plant health status and to monitor stress responses in large population screening, as it is extremely difficult to screen population for gas-exchange, especially in field conditions (Pask *et al*., 2012). However, Pask *et al*. (2012) noted that F_v_/F_m_ may not always be a reliable indicator of crop performance. While F_v_/F_m_ is often used as a proxy for photosynthetic performance in phenotyping, its link with gas exchange measurements is not yet fully established. Some studies have found positive correlations with mixed success for V_cmax_ (R^2^ = 0.02 - 0.13 and R ^2^ = 0.11 to 0.47 in Zhuang *et al*., 2020 and Mission *et al*., 2006, respectively), J (R^2^ = 0.09 – 0.28 in Zhuang *et al*., 2020) and A_max_ (R^2^ = 0.52, Ishida *et al*., 2014). In the present study, the cultivar models performance for gas-exchange (V_cmax_, J, g_m_, k_c_^c^, R_photoresp_, A_max_) and biochemical parameters (V_i_, V_t_) with F_v_F_m_Pc ranged from 0.57 to 0.85 for Cadenza and are nearly zero for Bobwhite due to its senescence behaviour. The same would have been expected at the whole plant scale (F_v_/F_m_Sc). However, the R ^2^_adj_ range of the two cultivars was between 0.32 to 0.75. Whilst results showed no significant differences between dates, a decreasing trend was observed, leading to an increase in model performances for Bobwhite at the whole plant scale. Despite the contrasting results between the two sets of measurements, the Interaction models gave similar performances, ranging from 0.28 to 0.66. This suggests that F_v_/F_m_ is a reliable indicator of photosynthetic performance at a seasonal scale, regardless of the measurement environment. Using F_v_/F_m_ as a predictor of the chlorophyll content (Chl a, b and SPAD) or the protein content variables (TSP, Rubisco) gave low to high performances depending on the cultivar (*i.e.*, senescence behaviour) and the scale of the measurement. These findings are consistent with the previous literature, which has reported high correlations between Fv/Fm and these variables accross different stages of development (Peng *et al*., 2017, Hailemichael *et al*., 2016, Jia *et al*., 2019) and/or under various nitrogen or irrigation treatments (Jia *et al*., 2019, Zarco-Tejada *et al*., 2013).

The PSII operating efficiency (Φ(II)), also known as quantum efficiency, provides an overall measurement of the efficiency of the PSII reaction centre in light conditions (Genty *et al*., 1989). This parameter indicates the proportion of light absorbed by chlorophyll associated with PSII that is used in photochemistry, and is strongly correlated with net CO2 assimilation (McAusland *et al*., 2013). It is also affected by the degradation of the photosystem during senescence as demonstrared in Mu *et al*. (2010) and Dai *et al*. (2004) where Φ(II) began to decrease significantly after 15 DPA. Similarly, Kong *et al*. (2015) observed a significant decrease in Φ(II) around the same time period. In the present study significant differences in Φ(II)_532_, Φ(II)FM and Φ(II)HP for Cadenza were observed after 14 DPA, consistent with previous research (Fig. 3). However, variations between dates were not uniform across instruments. No significant difference was observed between dates for Cadenza and Bobwhite with Φ(II)FP, as well as Φ(II)FM for Bobwhite. For Φ(II)HP, values reached their peak at 7 DPA and were significantly different at 28 DPA, while for Φ(II)_532_, the highest values were at booting and no significant differences were observed from anthesis to 28 DPA. The present study showed that Φ(II) under moderate light is a good estimator for carbon assimilation related parameters measuredeitherbygas-exchange(V_cmax_, J, g_m_, k_c_^c^, R_photoresp_, A_max_) or derived from biochemical analysis (V_i_ and V_t_). Except for k_c_^c^, the model performances were better for Cadenza than for Bobwhite, possibly due to the absence of significant differences between dates mentioned above. High and positive correlations between Φ(II) and V_cmax_, J, A_sat_ and g_m_ were also found in Sun *et al*. (2014), with variation caused by contrasted levels of ozone. Almaida *et al*. (2021) also observed positive correlations with A_max_ and V_cmax_ resulting from genetic variability of the sugar cane cultivars and wild types. Using Φ(II) as a predictor of the chlorophyll content will give reasonable results as long as both variables behave similarly during senescence.

Non-photochemical quenching (NPQ) is one of the three main mechanisms for dissipating the energy of the light absorbed by the chlorophyll pigments. NPQ competed with the photochemistry and the re-emission of the energy as a photon of fluorescence (Porcar-Castell *et al*., 2014). NPQ is a photoprotective mechanism removing the excess excitation energy within the chlorophyll-containing complexes (Murchie and Lawson, 2013). However, contradictory results have been reported in the literature regarding changes of NPQ after anthesis and during the senescence in wheat. Some authors observed an increase of NPQ during the period post anthesis period, with measurements performed either in the morning or at midday (Dai *et al*., 2004), and in different shading conditions (Mu *et al*., 2010). Xu *et al*. (2017) and Shao *et al*. (2013) observed an increase in NPQ peaking at 21 and 30 DPA, respectively, before decreasing. However, Kong *et al*. (2015) reported a significant decrease of NPQ measured on the flag leaves from anthesis to 32 DPA, and Peng *et al*. (2021) observed a decrease in NPQ between anthesis and maturity stages in wheat grown with various nitrogen inputs. The results of the present study showed a decreasing trend in Φ(NPQ)_1077_ from anthesis, becoming significant from 21 DPA for both cultivars compared to anthesis time. The model’s performances with the photosynthesis parameters derived from gas-exchange and from biochemical analysis were higher for Bobwhite (R ^2^_adj_ = 0.78 - 0.89) than Cadenza, indicating a better match between the senescence pattern of Φ(NPQ)_1077_ and those variables in Bobwhite. Negative correlations with net CO_2_ assimilation have been reported (Ishida *et al*., 2014, Zarco-Tejada *et al*., 2013), but no data concerning A_max_ or the other parameters derived from the A-C_i_ and A-PAR curves could be found in the literature to our knowledge. Negative correlations with Chl a content have been also reported in Ishida *et al*. (2014) and Zarco-Tejada *et al*. (2013), mainly driven by drought.

The handheld fluorescence devices were used as comparison with the fluorescence imagers, as well as with NDVI. While the use of NDVI to estimate F_v_/F_m_ and Φ(II) could be questionable, the results did not differ much from those reported in the literature, and low to moderate relationships were observed (Peng *et al*., 2017; Tan *et al*., 2012, Jia *et al*., 2019). The difference in performance between Bobwhite and Cadenza is attributed to their senescence behaviour, as discussed earlier. Usually, indices such as PRI, red-edge indices and fluorescence band indices are preferred to report variation in fluorescence. However, the hyperspectral image collection was done under non-optimal outdoor conditions, with variable ambient illumination between the end of September and the beginning of November in the UK. Consequently, the above-mentioned indices would not be reliable to report the variations in fluorescence, as the conditions between acquisition dates were too inconsistent.

The variation of F_v_/F_m_ measured by the two imagers reflected the variation of F_v_/F_m_ measured using the HandyPea for Cadenza. For Bobwhite, although no significance difference was observed between date for F_v_/F_m_Sc at the plant scale, the observed decreasing trend explained the relatively good correlation with the value collected with HandyPea. The measurements resulting from the Phenocenter did not show variations between time points, while the data collected with the HandyPea on the same leaves showed a significant decrease over time. The results are unexpected as the leaves were cut from the plants just before being imaged by the fluorescence of the Phenocenter, and no sensible explanation could be found.

The variation of Φ(NPQ)_1077_ was better reflected by the variation of F_v_/F_m_ than by Φ(II), in agreement with Kong et al (2015), where the decrease in NPQ appeared to be more concomitant with the decrease of F_v_/F_m_ than with the decrease of Φ(II). The authors also mentioned a regional accordance of imaging F_v_/F_m_ and NPQ across the leaves, concluding that this would indicate that NPQ is closely associated with the quantum efficiency of PSII photochemistry.

### Throughput consideration of the different approaches

Reynolds *et al*. (2012) noted that breeders are interested in reducing the size of their test populations to achieve the desired genotype at a reasonable probability and cost, even if the methods may be imperfect from a physiological and theoretical standpoint. In the case of photosynthetic traits, gas-exchange and biochemical analysis represent the gold standards but are also considered low throughput. The time required to perform an A-C_i_ followed by an A-PAR curve takes approximately 50-60 minutes per flag leaf for an experienced user in controlled conditions.

Biochemical analysis involves destructive sampling that also requires processing (freeze drying, grinding, etc) before quantification of TSP, Rubisco and enzymatic reactions. Handheld fluorescence instruments are used for pre-screening of populations for photosynthesis performance using either Φ(II) or F_v_/F_m_ as they are quite fast. Measuring Φ(II) with the Fluorpen is straightforward and takes only 10 seconds per leaf, potentially allowing for measurement of 360 leaves per hour, which is the fastest instrument to our knowledge. However, as shown in this study, the correlations with image fluorescence are weaker than those obtained with the HandyPea and the FMS2.

The advantages of HTP methods are the higher phenotyping capacity and, in some cases, the reduced necessity of human intervention. However, the throughput is highly dependent of the type of the technologies used. For example, in the case of the Phenocenter, an operator is required to program the execution, but the measurements are automatic and there is the possibility to perform an automatic loading of sample, pre inserted in the machine. The time frame for measuring F_v_/F_m_, followed by a light curve for Φ(II) and Φ(NPQ), on three flag leaves is around 40 minutes, including the 30 minutes of dark adaptation. The system should be able to host several boxes of three leaves for dark adaptation, allowing to measure 18 leaves per hour in total. The Field Scanalyzer requires an operator to program the platform, but the platform runs automatically over the full experiment once it is launched. The throughput of the fluorescence camera is 40 plots (or pots) per hour, but this includes the time it takes to move the camera between targets. The visible diameter of the flash used to induce fluorescence is 1.2 to 1.4 meters, which can also trigger a response to light in nearby plots. Therefore, the device must be carefully routed to measure each plot in complete dark adaptation, which decreases the camera’s throughput. The hyperspectral camera, set up in medium spatial resolution (Sadeghi-Tehran *et al*., 2021), allows collection of 26 scans per hours. As the area covered is about 1.8 x 1.5 m, the number of plots or pots scanned will vary depending on the size of targets. The two phenotyping platforms discussed in the previous paragraph offer significant advantages over gas-exchange measurements in terms of speed and reduced human intervention. However, their throughput is still quite low compared to handheld fluorescence instruments, which are commonly used as proxy of photosynthesis. Nonetheless, spectral data are still promising for fluorescence and photosynthetic traits. The Fraunhofer Line Depth (FLD) can be used to retrieve sun-induced fluorescence (Porcar-Castell *et al*., 2014) and the last decade have seen an increase of prediction of V_cmax_ and J using the full range of spectral data (Silva-Perez *et al*., 2018; Meacham-Hensold *et al*., 2019, Montes *et al*., 2022). Most of those developments have relied on single spectroradiometer, as they are much faster than the imager mounted on ground-based system. However, hyperspectral imaging with UAV provides data on large scale, enabling access to those traits at lower spatial resolution (Zarco-Tejada *et al*., 2013; Camino *et al*., 2018, 2019).

#### Conclusion

The present study showed that photosynthetic traits obtained either by gas-echange or biochemical analysis could be monitored and quantified susccesfully using imaging technologies as long as the senescence is driving the variability. Despite the contrasted behaviour between the two cultivars, Bobwhite and Cadenza, linear regression models which account for both the cultivar and the interaction effects, further improved the modelling of photosynthesis indicators. Future work will be necessary to evaluate the link between the image variables and gas-exchange during earlier stages of development. The results also showed the importance of prior knowledge of the stay green type of the cultivar used, as it can strongly impact the outcomes. It also emphasises the need to develop sensitive enough approaches to differentiate cosmetic from functional stay green genotypes that breeders can use for photosynthetic traits.

## Supplementary data

Th following supplementary data are available at the JXB online:

Fig. S1. Flag leaf holder for Phenocenter analysis

Fig. S2. Pots settings under the Field Scanalyzer

Fig. S3. Linear regressions between Fv/FmSc and selected variables

Fig. S4. Linear regressions between NDVI and selected variables

Fig. S5. Linear regressions between F_v_/F_m_Pc and selected variables

Fig. S6. Linear regressions between Φ(II)_532_ and selected variables

Fig. S7. Linear regressions between Φ(NPQ)_1077_ and selected variables

Table S1. Linear regressions parameters between images and A-C_i_ curves

Table S2. Linear regressions parameters between images and A-PAR curves

Table S3. Linear regressions parameters between images and biochemical variables

Table S4. Linear regressions parameters between images and handhled devices variables

Table S5. Mean and standard deviation for each variables per date and per cultivar

Dataset S1: Phenotypic data

## Acknowledgments

We thank Kirsty Hassal and Jess Evans for their help in disigning the experiment and their advice in the analysis of the data (Rothamsted Research), Fiona Gilzean for the access to the CE room and cabinet for growth of the plants.

## Author contributions

JPP and NV designed and performed the experiments and analysed data; TA performed the image acquisition at the PhenoCenter (CHAP), PST performed the image extraction and analysis from the Field Scanalyzer; DO and ECS peformed the biochemical analyses; NV and JPP wrote the manuscript; NV, JPP, TA, PST, DO, ECS and MJH revised and edited the manuscript; JPP, ECS and MJH enabled to secure funding for the project leading to this publication. All authors reviewed and approved the final manuscript.

## Data availability

All data supporting the findings of this study are available within the paper and within its supplementary data published online.

## Abbreviations

V_cmax_: maximum carboxylation rate
J: CO_2_-saturated electron transport rate
g_m_: mesophyll conductance
R_photoresp_: photorespiration
k_c_^c^: carboxylation efficiency
A_max_: maximum net assimilation rate
TSP: total soluble protein content
Chl a and b: chlorophyll and b content
Rubisco: Rubisco content
V_i_: initial Rubisco activity
V_t_: total Rubisco activity
Φ(II): PSII operating efficiency or quantum yield
F_v_/F_m_: maximum efficiency of the photosystem II
Φ(NPQ): non-photochemical quenching
NDVI: normalised difference vegetation index
DPA: Days post anthesis.

