## Supplementary Figures for "A multiscale approach to investigate fluorescence and NDVI imaging as proxy of photosynthetic traits in wheat"

Figure S1. Device for flag leaf holding for Phenocenter analysis (A); Device inserted in the Phenocenter for fluorescence analysis (B).

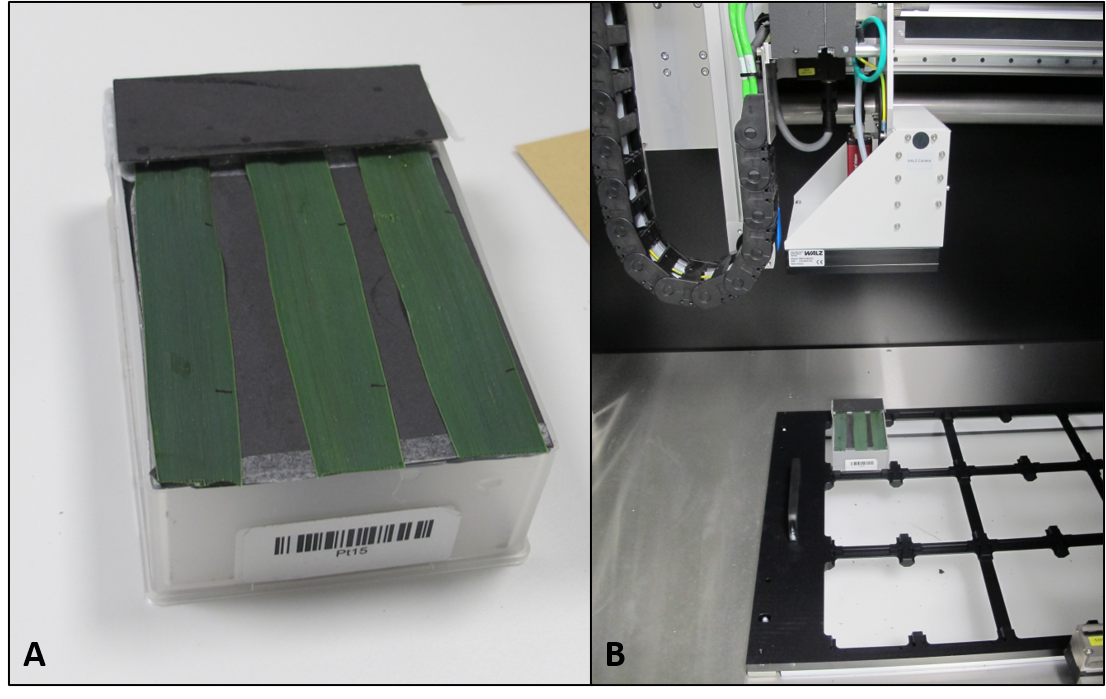

Figure S2. Pots allocated to the Field Scanalyzer analysis.

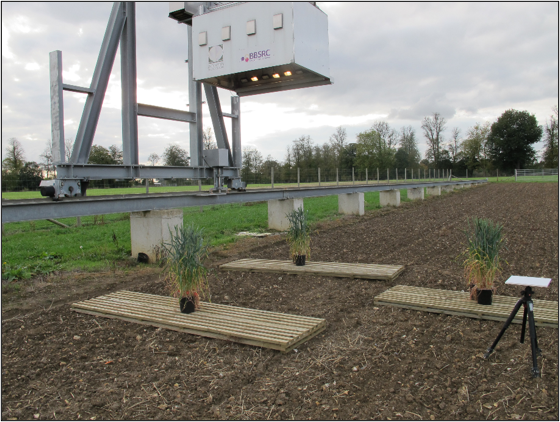

Figure S3. Linear regressions between chlorophyll fluorescence (F_v_/F_m_Sc) measured by the Field Scanalyzer and other measured parameters.

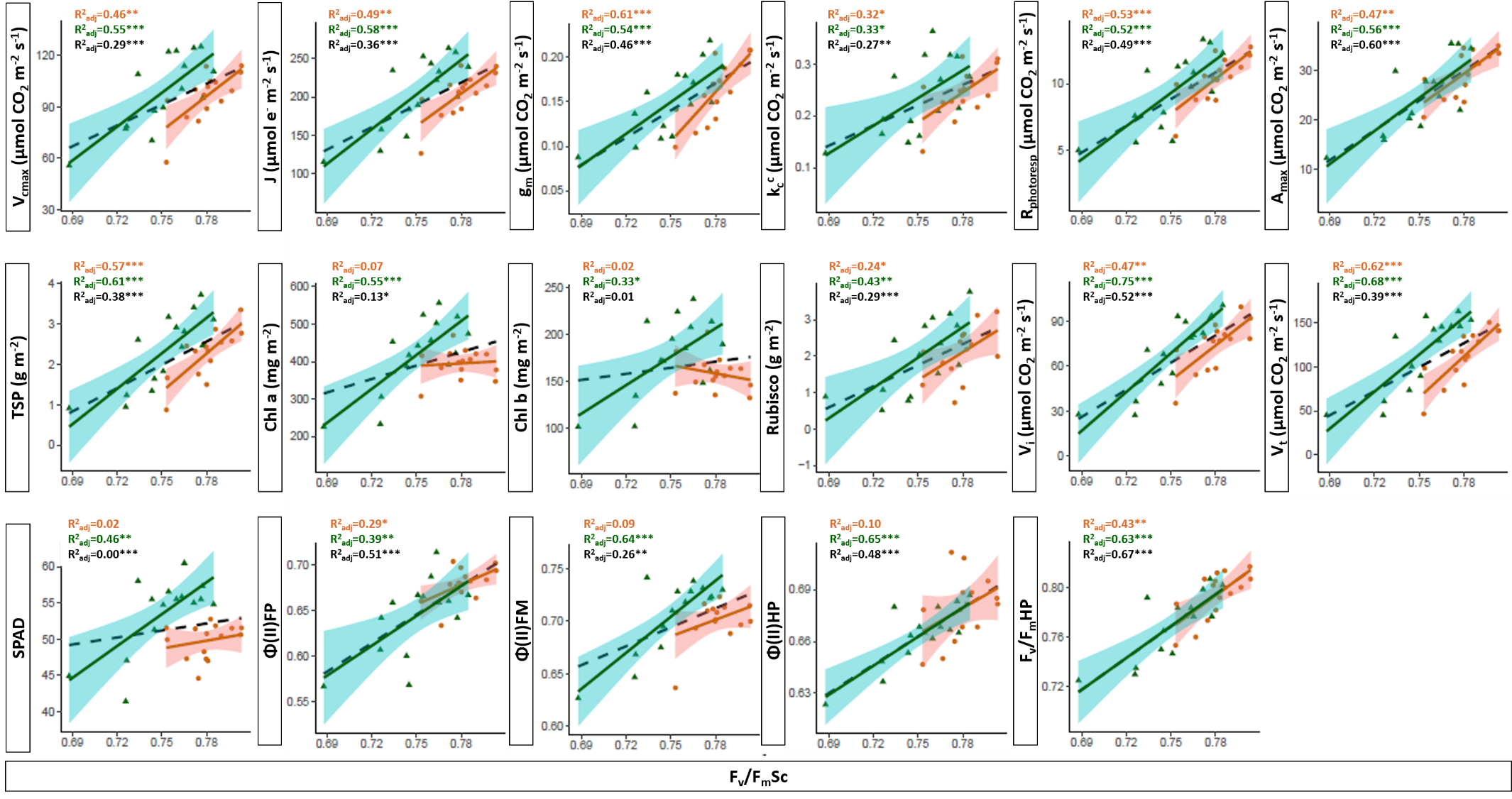
Bobwhite, orange circles and solid lines; Cadenza, green triangles and solid line. The dashed black lines is for the generalist model. R^2^, coefficient of determination of the regression. ***, *p* < 0.001; **, *p* < 0.01; *, *p* < 0.05.

Figure S4. Linear regressions between normalized difference vegetation index (NDVI) measured by the Field Scanalyzer and other measured parameters.

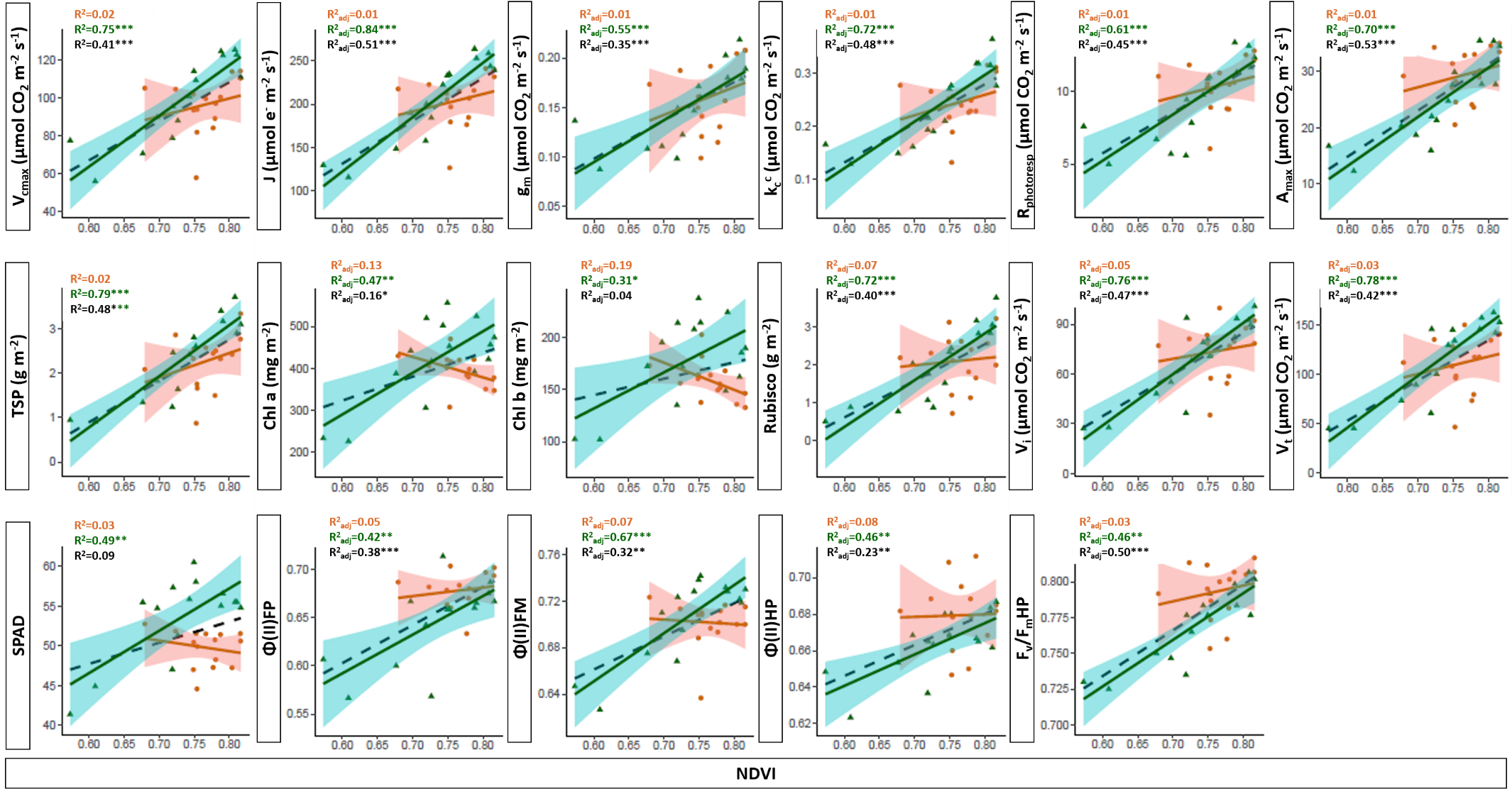
Bobwhite, orange circles and solid lines; Cadenza, green triangles and solid line. The dashed black lines is for the generalist model. R^2^, coefficient of determination of the regression. ***, *p* < 0.001; **, *p* < 0.01; *, *p* < 0.05.

Figure S5. Linear regressions between chlorophyll fluorescence (F_v_/F_m_Pc) measured by the Phenocenter and other measured parameters.

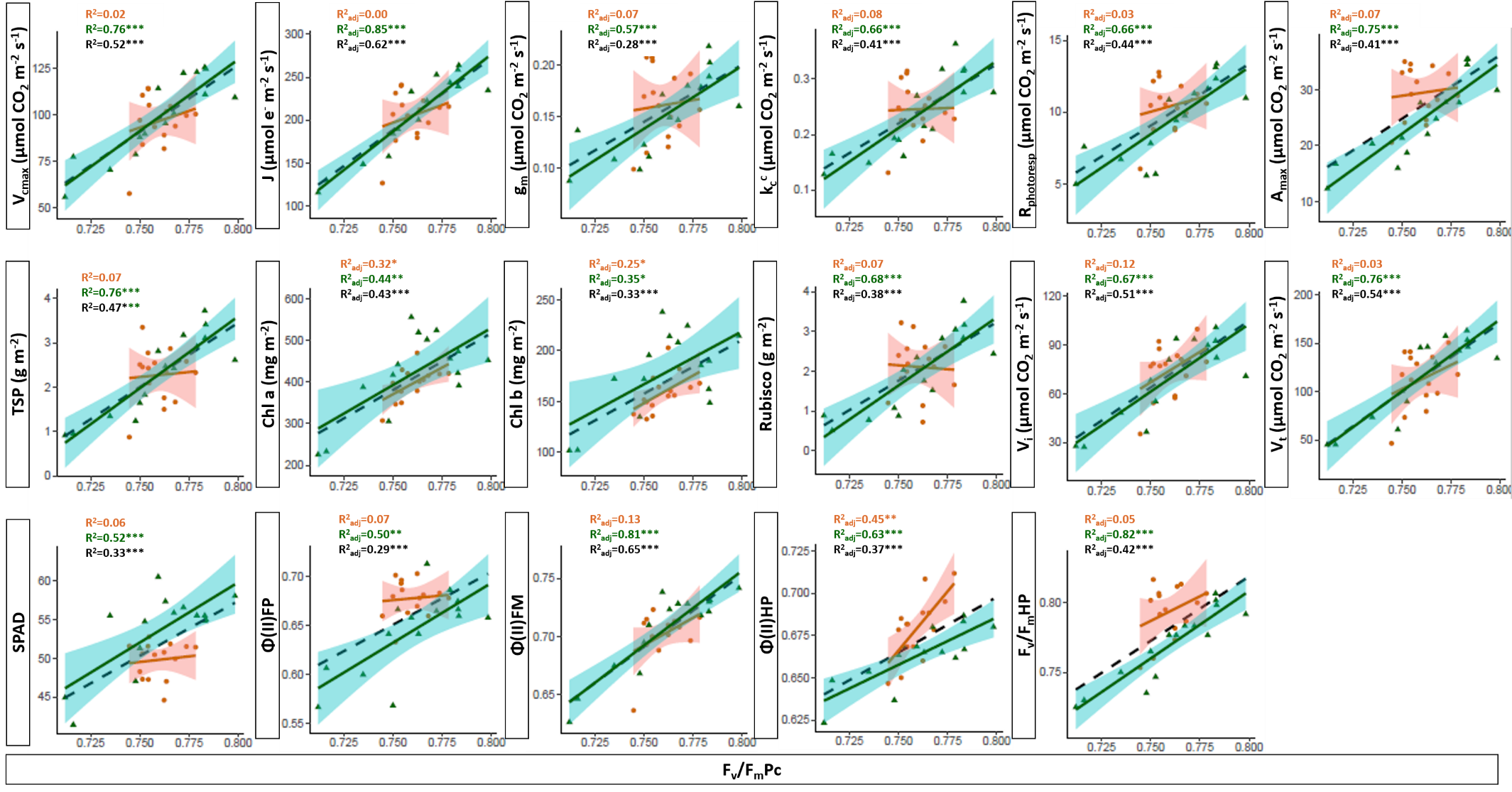
Bobwhite, orange circles and solid lines; Cadenza, green triangles solid line. The dashed black lines is for the generalist model. R^2^, coefficient of determination of the regression. ***, *p* < 0.001; **, *p* < 0.01; *, *p* < 0.05.

Figure S6. Linear regressions between photosystem II quantum efficiency (Φ(II)_532_) measured by the Phenocenter and other measured parameters.

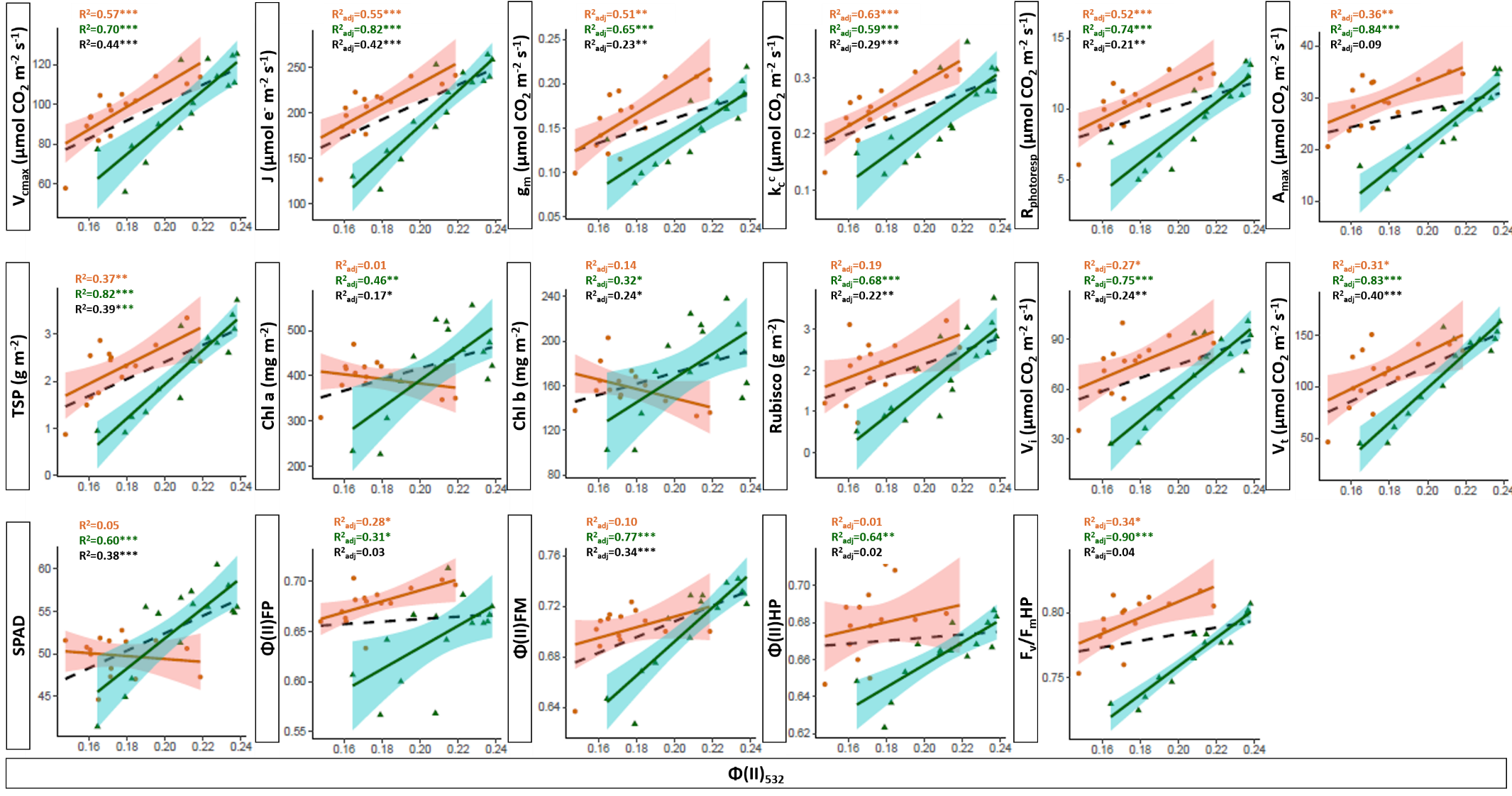
Bobwhite, orange circles and solid lines; Cadenza, green triangles and solid line. The dashed black lines is for the generalist model. R^2^, coefficient of determination of the regression. ***, *p* < 0.001; **, *p* < 0.01; *, *p* < 0.05. Φ(II)_532_, photosystem II quantum efficiency at light level of 532 µmols photons m^-2^ s^-1^.

Figure S7. Linear regressions between non-photochemical quenching (Φ(NPQ)_1077_) measured by the Phenocenter and other measured parameters.

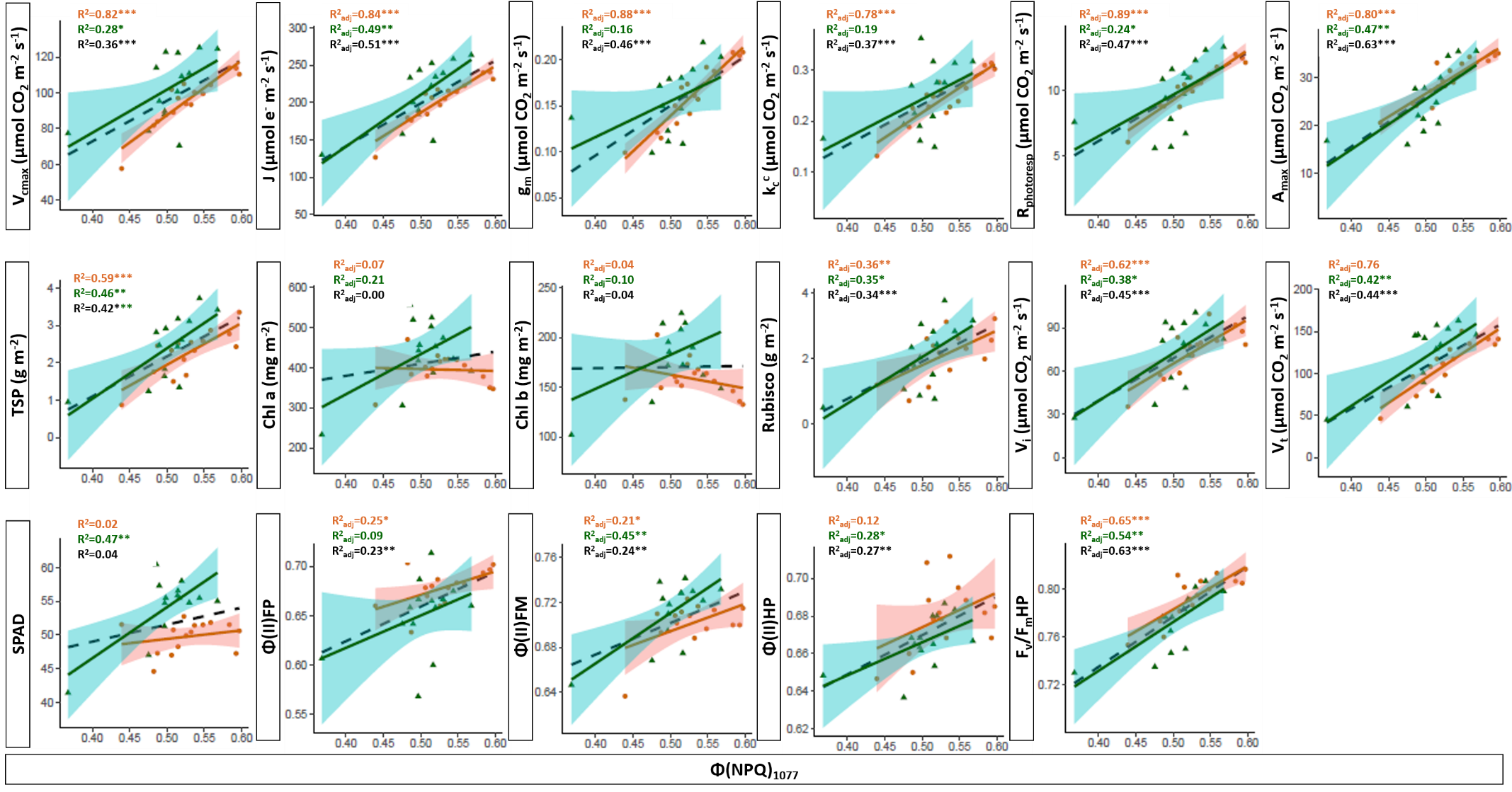
Bobwhite, orange circles and solid lines; Cadenza, green triangles and solid line. The dashed black lines is for the generalist model. R^2^, coefficient of determination of the regression. ***, *p* < 0.001; **, *p* < 0.01; *, *p* < 0.05. Φ(NPQ)_1077_, non-photochemical quenching at light level of 1077 µmols photons m^-2^ s
