## Supplementary Tables for "A multiscale approach to investigate fluorescence and NDVI imaging as proxy of photosynthetic traits in wheat"

Supplementary Table 1. General parameters of the linear regressions between high-throughput methods and parameters originated from A-C_i_ curves

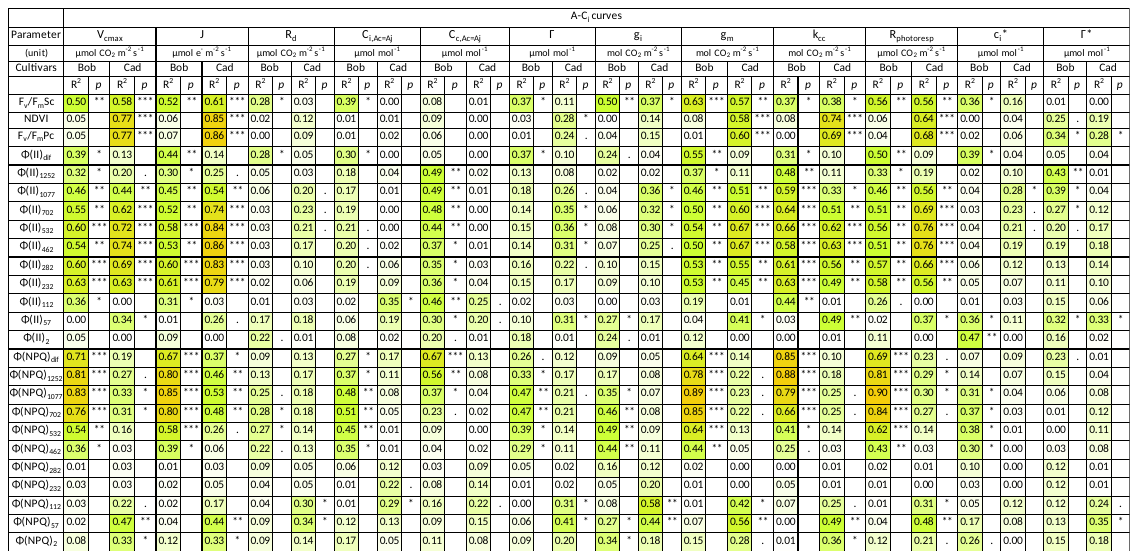

*p*, p-value of the regression; R^2^, coefficient of determination of the regression. ***, *p* < 0.001; **, *p* < 0.01; *, *p* < 0.05.

Supplementary Table 2. General parameters of the linear regressions between high-throughput methods and parameters originated from A-PAR curves

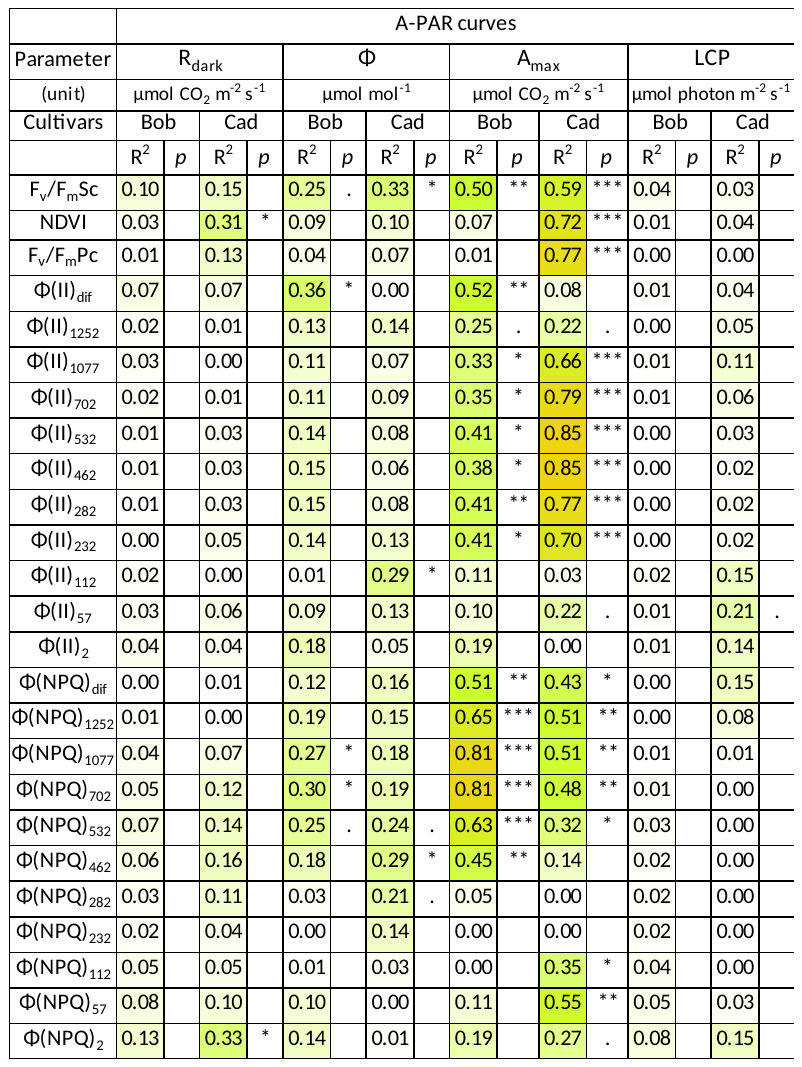

*p*, p-value of the regression; R^2^, coefficient of determination of the regression. ***, *p* < 0.001; **, *p* < 0.01; *, *p* < 0.05.

Supplementary Table 3. General parameters of the linear regressions between high-throughput methods and parameters originated from biochemical analysis

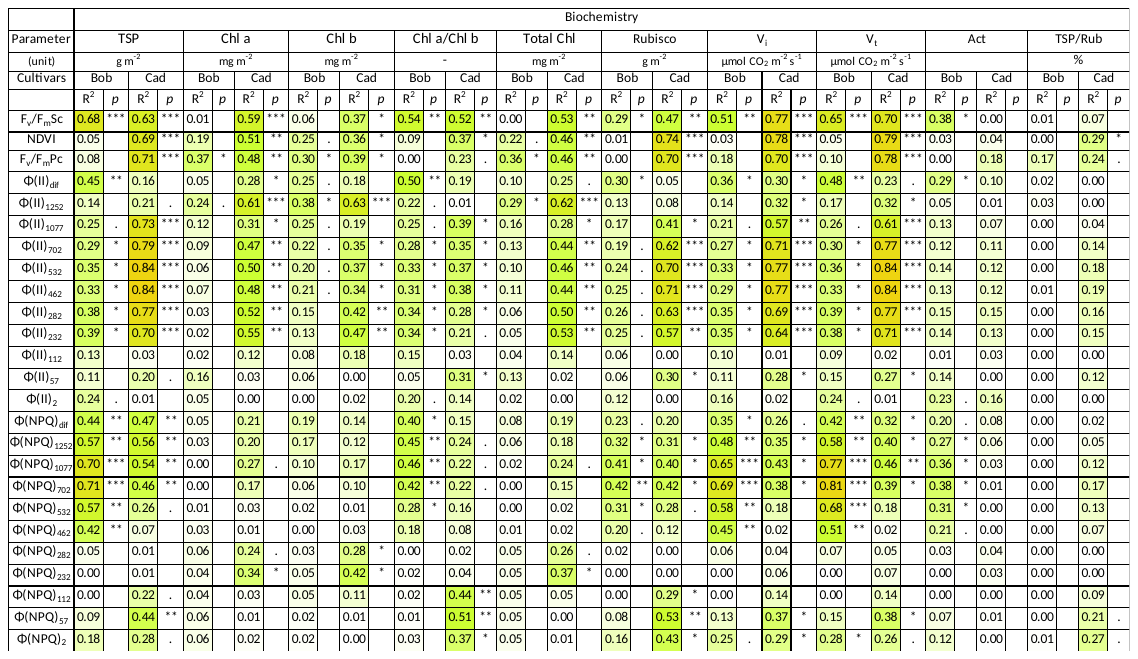

*p*, p-value of the regression; R^2^, coefficient of determination of the regression. ***, *p* < 0.001; **, *p* < 0.01; *, *p* < 0.05.

Supplementary Table 4. General parameters of the linear regressions between high-throughput methods and parameters originated from chlorophyll and fluorescence evaluations

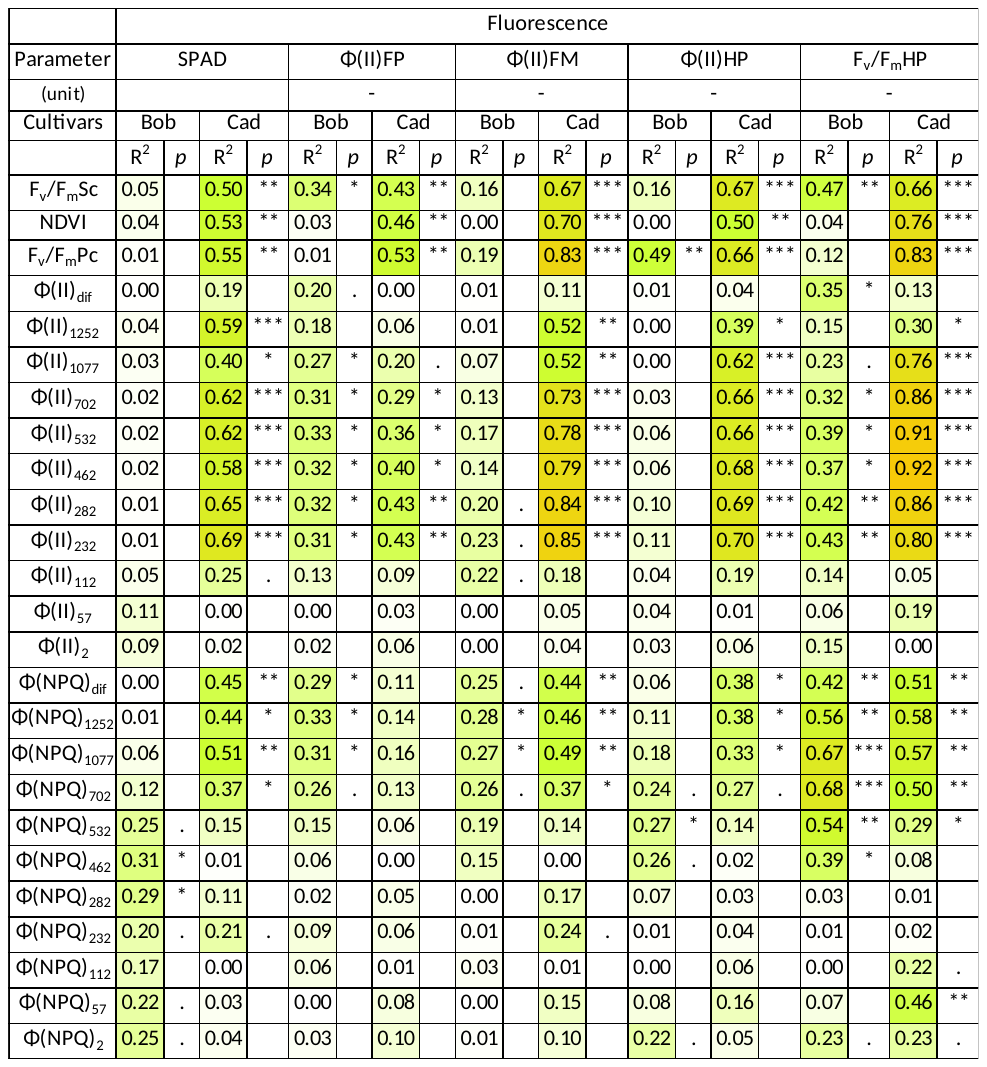

*p*, p-value of the regression; R^2^, coefficient of determination of the regression. ***, *p* < 0.001; **, *p* < 0.01; *, *p* < 0.05.

| Time Point | Z4.5 | | | | Z6.5 | | | | Z6.5+7 | | | | Z6.5+14 | | | | Z6.5+21 | | | |
| --- | --- | --- | --- | --- | --- | --- | --- | --- | --- | --- | --- | --- | --- | --- | --- | --- | --- | --- | --- | --- |
| Cultivar | Bobwhite | | Cadenza | | Bobwhite | | Cadenza | | Bobwhite | | Cadenza | | Bobwhite | | Cadenza | | Bobwhite | | Cadenza | |
| GS | Z4.5 | | Z4.5 | | Z6.5 | | Z6.5 | | Z7.3 | | Z7.1 | | Z7.7 | | Z7.9 | | Z8.3 | | Z8.3 | |
| DAS | 43 | | 54 | | 48 | | 63 | | 55 | | 70 | | 62 | | 77 | | 69 | | 84 | |
|  | Mean | Std | Mean | Std | Mean | Std | Mean | Std | Mean | Std | Mean | Std | Mean | Std | Mean | Std | Mean | Std | Mean | Std |
| V_cmax_ | 112.8 | 2.075 | 120.2 | 8.143 | 99.15 | 5.527 | 118.1 | 7.708 | 99.70 | 2.460 | 103.3 | 9.634 | 95.98 | 8.217 | 82.70 | 10.61 | 74.51 | 14.58 | 70.64 | 12.86 |
| J | 237.5 | 5.266 | 253.9 | 12.82 | 214.2 | 8.694 | 243.7 | 9.103 | 211.6 | 4.552 | 218.5 | 16.84 | 199.7 | 16.60 | 174.2 | 22.25 | 160.9 | 29.66 | 134.4 | 21.35 |
| g_m_ | 0.207 | 0.002 | 0.203 | 0.015 | 0.180 | 0.017 | 0.173 | 0.011 | 0.159 | 0.010 | 0.156 | 0.014 | 0.148 | 0.022 | 0.114 | 0.008 | 0.111 | 0.011 | 0.108 | 0.026 |
| k_c_^c^ | 0.309 | 0.006 | 0.303 | 0.023 | 0.240 | 0.024 | 0.319 | 0.044 | 0.243 | 0.013 | 0.231 | 0.033 | 0.254 | 0.024 | 0.166 | 0.021 | 0.182 | 0.048 | 0.162 | 0.033 |
| R_photoresp_ | 12.48 | 0.313 | 12.93 | 0.540 | 11.19 | 0.640 | 11.30 | 0.326 | 10.45 | 0.156 | 10.01 | 0.756 | 9.911 | 1.132 | 6.738 | 1.071 | 7.910 | 1.614 | 6.048 | 1.364 |
| A_max_ | 34.37 | 0.795 | 35.14 | 0.562 | 32.86 | 1.442 | 28.46 | 1.255 | 29.72 | 3.027 | 24.80 | 2.818 | 27.02 | 2.973 | 20.13 | 1.329 | 23.07 | 2.209 | 14.99 | 2.381 |
| TSP | 2.847 | 0.464 | 3.410 | 0.307 | 2.663 | 0.176 | 2.897 | 0.282 | 2.387 | 0.104 | 2.566 | 0.212 | 1.750 | 0.304 | 1.603 | 0.244 | 1.694 | 0.791 | 1.031 | 0.178 |
| Chl a | 358.4 | 16.82 | 429.2 | 41.52 | 415.5 | 8.176 | 478.0 | 40.44 | 406.7 | 12.59 | 526.2 | 27.60 | 408.2 | 26.03 | 415.7 | 27.26 | 387.1 | 81.44 | 254.9 | 44.04 |
| Chl b | 138.4 | 7.122 | 166.9 | 20.96 | 161.6 | 4.601 | 208.2 | 20.17 | 160.1 | 8.160 | 220.1 | 15.81 | 170.4 | 13.49 | 179.9 | 13.39 | 163.4 | 34.72 | 113.1 | 18.96 |
| Rubisco | 2.589 | 0.613 | 3.251 | 0.473 | 2.681 | 0.410 | 2.767 | 0.309 | 2.219 | 0.498 | 1.873 | 0.424 | 1.811 | 0.588 | 1.227 | 0.698 | 1.246 | 0.548 | 0.822 | 0.286 |
| Vi | 86.09 | 6.991 | 91.76 | 9.315 | 86.41 | 11.62 | 84.67 | 12.13 | 79.98 | 3.249 | 84.62 | 8.031 | 68.92 | 9.406 | 57.15 | 10.06 | 49.03 | 12.00 | 30.48 | 5.187 |
| Vt | 139.0 | 3.675 | 154.0 | 8.536 | 138.2 | 11.13 | 144.6 | 11.81 | 114.4 | 5.435 | 140.3 | 9.326 | 96.60 | 16.37 | 87.79 | 13.62 | 71.92 | 24.93 | 50.17 | 8.993 |
| SPAD | 49.81 | 2.252 | 55.12 | 0.349 | 51.27 | 0.731 | 56.71 | 1.264 | 48.91 | 2.274 | 57.89 | 2.375 | 51.16 | 1.456 | 53.84 | 2.230 | 47.82 | 3.513 | 44.46 | 2.844 |
| ɸ(II)FP | 0.697 | 0.004 | 0.667 | 0.008 | 0.676 | 0.011 | 0.670 | 0.015 | 0.679 | 0.001 | 0.671 | 0.038 | 0.672 | 0.013 | 0.612 | 0.050 | 0.666 | 0.035 | 0.605 | 0.038 |
| ɸ(II)FM | 0.705 | 0.009 | 0.728 | 0.005 | 0.699 | 0.013 | 0.729 | 0.012 | 0.712 | 0.004 | 0.730 | 0.008 | 0.711 | 0.011 | 0.693 | 0.018 | 0.680 | 0.038 | 0.647 | 0.021 |
| ɸ(II)HP | 0.678 | 0.009 | 0.679 | 0.011 | 0.684 | 0.014 | 0.669 | 0.010 | 0.699 | 0.018 | 0.671 | 0.008 | 0.683 | 0.005 | 0.662 | 0.008 | 0.652 | 0.007 | 0.636 | 0.013 |
| F_v_/F_m_HP | 0.809 | 0.006 | 0.802 | 0.004 | 0.803 | 0.009 | 0.784 | 0.008 | 0.807 | 0.005 | 0.779 | 0.004 | 0.787 | 0.005 | 0.754 | 0.010 | 0.762 | 0.010 | 0.730 | 0.005 |
| F_v_/F_m_Sc | 0.795 | 0.014 | 0.778 | 0.007 | 0.791 | 0.006 | 0.749 | 0.014 | 0.777 | 0.004 | 0.769 | 0.008 | 0.772 | 0.016 | 0.746 | 0.004 | 0.765 | 0.011 | 0.713 | 0.022 |
| NDVI | 0.812 | 0.007 | 0.804 | 0.014 | 0.746 | 0.022 | 0.785 | 0.030 | 0.772 | 0.021 | 0.737 | 0.016 | 0.738 | 0.052 | 0.700 | 0.025 | 0.761 | 0.014 | 0.633 | 0.076 |
| F_v_/F_m_Pc | 0.753 | 0.002 | 0.783 | 0.000 | 0.766 | 0.008 | 0.783 | 0.014 | 0.764 | 0.014 | 0.763 | 0.004 | 0.762 | 0.007 | 0.746 | 0.010 | 0.753 | 0.009 | 0.725 | 0.020 |
| ɸ(II)_532_ | 0.208 | 0.012 | 0.237 | 0.001 | 0.166 | 0.005 | 0.221 | 0.013 | 0.179 | 0.006 | 0.219 | 0.008 | 0.166 | 0.010 | 0.198 | 0.009 | 0.161 | 0.012 | 0.175 | 0.010 |
| ɸ(NPQ)_1077_ | 0.591 | 0.007 | 0.547 | 0.019 | 0.546 | 0.013 | 0.511 | 0.011 | 0.519 | 0.016 | 0.496 | 0.015 | 0.519 | 0.009 | 0.503 | 0.011 | 0.470 | 0.026 | 0.422 | 0.076 |

Supplementary Table 5: Mean and standard deviation of the 3 replication of each considered variables per date and per cultivar.
